# Nature protection across countries: Do size and power matter?

**DOI:** 10.1101/861971

**Authors:** Germán Baldi

## Abstract

Protected areas are one of the most effective tools for nature conservation. Consequently, almost all countries have agreed to set increasingly demanding goals for the expansion of their protected area systems. However, there is a large disparity among countries, and research on the cultural drivers of differences remains quite unexplored. Here, we explore the relationship between protected extent and a limited spectrum of socio-economic characteristics, making focus on size and power features. Protected areas under strict conservation categories (I to IV, IUCN) were considered for 195 countries, and relationships were modeled by means of LOESS regressions, violin plots, and a random forest ensemble learning method. Larger and more powerful countries (in terms of land area, gross domestic product, or military expenditure) protect less and in relatively smaller units than smaller and less powerful countries. Out of the twenty most extensive countries of the world, only two exceed 10% of protection. This situation is problematic since an effective growth of the global protected area network depends on the willingness of larger and more powerful countries. We propose different hypotheses *a posteriori* that explain the role of size and power driving protection. These hypotheses involve direct mechanisms (e.g., the persuasive capacity of large countries) or mechanisms that mediate the interactions of some others (e.g., tourism contribution to GDP and insularity). Independently of mechanisms, our results emphasize the conservation responsibilities of large and powerful countries and contribute to envision conservation scenarios in the face of changes in the number and size of countries.

## 1. Introduction

The physical, biological and cultural assets of the planet are unequally divided among more than 200 sovereign countries and dependencies of different legal character. The six largest countries of the world occupy 45% of the land area excluding Antarctica, while the smallest hundred occupy only 2.5% (Gini index, G = 0.80, Table 1; data sources are depicted in Table 2). Moreover, ten countries exceeding 100 M inhabitants, encompass 60% of the world population, while there are 115 countries with less than 1 M (G = 0.81). This conjunction of conditions clearly implies differential access and appropriation of natural resources by humans, which is reflected and magnified in the gross domestic product (G = 0.87) and in the military expenditure (G = 0.90). Perhaps less obvious is that these inequalities together imply different degrees of responsibility on the part of the administrations of countries in the long-term conservation of natural and cultural assets. This makes global sustainability hard to achieve considering that it should be a joint effort which exceeds current and future political borders.

**Table 1.** Inequality in the distribution of variables related to the size and power of a country, following Crowards (2002b, a) and Arvanitidis and Kollias (2016).

| Variable | Minimum | Median | Maximum | Inequality<br>(Gini index) |
| --- | --- | --- | --- | --- |
| Land area (k km<sup>2</sup>) | 0.01 x 10<sup>-3</sup> | 566 | 16,953 | 0.80 |
| Gross domestic<br>product (M US<br>Dollars) | 10 | 2.7 x 10<sup>4</sup> | 1.6 x 10<sup>7</sup> | 0.87 |
| Population size (M<br>inh) | 8 x 10<sup>-4</sup> | 7.2 | 1,338 | 0.81 |
| Military<br>expenditure (M US<br>Dollars) | 0 | 412 | 6.5 x 10<sup>5</sup> | 0.90 |

**Table 2.** List of independent variables associated with the extent of the protected area by country. Variables #1 to #6 are related to the two intertwined concepts of size and power following Crowards (2002b) and Arvanitidis and Kollias (2016). Variables #7 to #16 are used to contextualize size and power results.

| # | Variable | Units | Description |
| --- | --- | --- | --- |
| 1 | **Crowards' country size** | Categorical | Crowards (2002b, a) classification of 179 independent countries into five categories, according to their area, GDP and population. The categorization of unclassified countries (e.g., Australia, Canada) were completed following his classification scheme |
| 2 | **Land area** | km<sup>2</sup> | Abundance and diversity of natural assets. From Natural Earth (2017) |
| 3 | **Gross domestic product (GDP)** | US Dollars | Economic size or development of a country and, therefore, political strength. From the World Bank (2018) average values 2007-2016 |
| 4 | **Population size** | Inh | Stock of human capital and size of the domestic market. From Natural Earth (2017) |
| 5 | **Military expenditure** | US Dollars | Capability to dissuade and/or coerce external entities/policies to either protect and/or advance national interests at national to global levels. Average values 2007-2016. From the World Bank (2018). Missing cases (e.g., Comoros), from the GlobalSecurity.org (2011). Other measures of military power has been proposed (Singer 1988, Beckley 2018), but due to redundancies in results, they are only presented on Figure S2, Appendix 1 |
| 6 | **External possessions** | categorical | Similar to #5, but enabling countries to influence policies abroad. Included countries (†) posses overseas military bases or govern overseas territories, territories with ethnic minorities and listed as "Non Self Governing Territories" according to the 2018 Session of the United Nations (e.g., American Samoa), "Special member state territories" of the European Union (e.g., Canary Is.) or other categories (e.g., Easter Is.) |
| 7 | **GDP per capita** | US Dollars * inh<sup>-1</sup> | Affluence of population. From the World Bank (2018) average values 2007-2016 |
| 8 | **Income inequality** | unitless | Estimate of inequality in equivalized (square root scale) household disposable income (post-tax, post-transfer) from Solt (2016). Average Gini index 2007-2016. For Andorra, The Bahamas, Cuba, North Korea, Saudi Arabia, and Singapore, from the IMUNA country profiles (2018) |
| 9 | **Education** | unitless | From the Human Development Index ranks. Calculated using mean years of schooling and expected years of schooling. Average values 2005-2015 (UNDP 2015) |
| 10 | **Population density** | inh * km<sup>-2</sup> | Pressure of population per area for natural resources and space. From Natural Earth (2017) |
| 11 | **Globalization** | unitless | Economic, social and political globalization, which includes data on economic flows and restrictions, information flow, and cultural proximity. From the KOF Swiss Economic Institute average values 2006-2015, overall values (Gygli et al. 2018) |
| 12 | **Governance** | unitless | Considering voice and accountability, political stability and absence of violence, government effectiveness, regulatory quality, rule of law, and control of corruption (Kaufmann et al. 2010). From the World Bank (2018) average values 2007-2016 |
|  |  |  | considering the average of the six governance components |
| 13 | **Tourism contribution to GDP** | % | Relative importance of this economic activity, given that nature protection attracts visitors interested in remarkable natural or cultural landscapes, and tourism drive protection to preserve this quality. From the WTTC (2017) data base. 2017 values. |
| 14 | **Age of the PA network** | Y | Number of years since the creation of the first protected area. From UNEP-WCMC (2018) |
| 15 | **Continental or insular** | categorical | If the country is mostly continental or insular (e.g., Malaysia is considered an insular country, as 60% of its territory is in Borneo Is.) |
| 16 | **Continent** | categorical | Following the six fold model (Lewis and Wigen 1997)<sup>‡</sup> |

† Australia, Chile, China, Denmark, Finland, France, Israel, Morocco, Norway, New Zealand, Portugal, Russia, Spain, The Netherlands, Turkey, United Kingdom of Great Britain and Northern Ireland, and United States of America.
‡ Russia is included in Europe.

The countries with the highest level of wealth were the first to formalize conservation policies under different agreements. With the sanction of the Yellowstone National Park Act in 1872, the United States of America installed the modern concept of protected area at a global level (Watson et al. 2014). This legal title entails the effective preservation of large tracts of land, which are later made available for public use. In the late 19th and early 20th centuries, other sovereign or colonial governments quickly adopted this model, such as Canada, Chile or South Africa (Szafer 1973, McNeely et al. 1994, Watson et al. 2014). The 20th century brought relevant geopolitical changes, such as the independence of colonial territories and the split of former territories after both World-Wars. In face of these changes, academic circles in Western Europe anticipated a degazettement of protected areas in Africa, Asia and Oceania. Conversely, these new countries actively promoted the creation of protected areas, as conservation was no longer exclusive to powerful countries (Frank et al. 2000, Fairbrother 2012). In a few years, peripheral and young countries, such as the Central African Republic or Belize established numerous protected areas (Radeloff et al. 2013). As of July 2018, there are around 240,000 terrestrial protected areas, which occupy a land area of 20.2 M km^2^ or 14.9% of the world land surface excluding Antarctica (UNEP-WCMC 2018). Furthermore, almost all countries in 2010 negotiated that at least 17% of terrestrial areas needed to be included within protected networks by 2020 (first clause of the Aichi Biodiversity Target 11, SCBD 2010). Without considering the city-states, only a few territories with diverse socio-economic conditions are completely devoid of protected areas, or their conservation systems have scarcely been developed (e.g., Haiti, Turkey) (McNeely et al. 1994).

Protected areas arise from a complex interplay of motivations related to perceived societal benefits, like the early preservation of iconic landscape features, or the late widespread agreement on the importance of maintaining nature and biodiversity (Pressey 1994, Watson et al. 2014, Baldi et al. 2017). However, these motivations act as underlying forces, being the process of conforming protected areas driven by direct human-related or cultural drivers (hereafter, cultural drivers). These drivers can be associated with the economic or political context of the country, its social organization, and prevailing moral rules (Baldi et al. 2017). There is currently a broad consensus about the need to increase the extent under protection in order to preserve the structure and functioning of nature (Rodrigues et al. 2004, Watson et al. 2014). That is why a joint analysis of these <u>drivers</u> would identify the conditions that may facilitate or boost the creation of protected areas, as well as the conservation debts maintained by countries that have favorable conditions, but for which conservation has played a minor role in political agendas.

Research on cultural drivers has not yet been examined or discussed thoroughly, and literature on this topic is mostly based on qualitative analyses focused on a few features. For example, several narratives have evidenced the importance of individual actors (e.g., heads of state, naturalists and scientists) and political actions driving the deployment of protected areas (Wells and Williams 1998, Pauchard and Villarroel 2002, Castañeda Rincón 2006, Ouyang et al. 2013, Leal 2017). Quantitative studies are scarcer. For example, Marinaro et al. (2012) showed that, in Argentina, the totalitarian administrations of the first half of the 20th century established protected areas of large size. Meanwhile, subsequent democratic administrations made focus on the diversification of the protected network, with the inclusion of many areas of small size only when significant economic surpluses were available.

Studies about cultural drivers becomes even less at the global level (Table 3). In their seminal assessment, Frank et al. (2000) showed that the links of countries to world society (e.g., the signature of environmental treaties) were strong drivers of the extent of protected areas. Later, Upton et al. (2008) found few significant relationships between poverty indicators and the extent of the protected area. They suggested that inconclusive associations could be attributed to the joint effect of two factors. On the one hand, local society pressures and economic capacity of more affluent countries aimed at setting aside land for conservation. On the other hand, international agendas and foreign investment in less affluent countries contributed to land conservation. McDonald and Boucher (2011) found that more affluent countries achieved a larger extent of protection in the middle of the 20th century, while currently, less affluent/developing and more affluent/developed countries protect their territories at similar rates. Kashwan (2017) found that protected extent depended mainly on the interaction between democratic strength and economic inequality. Protected areas tended to emerge under undemocratic settings with high inequality, or under democratic settings with low inequality. Finally, Baynham-Herd et al. (2018) assessed the engagement and investment in the environment and pro-environmental behavior and found that governance was the most relevant driver of protection, with a minor contribution of the level of globalization as well.

**Table 3.**
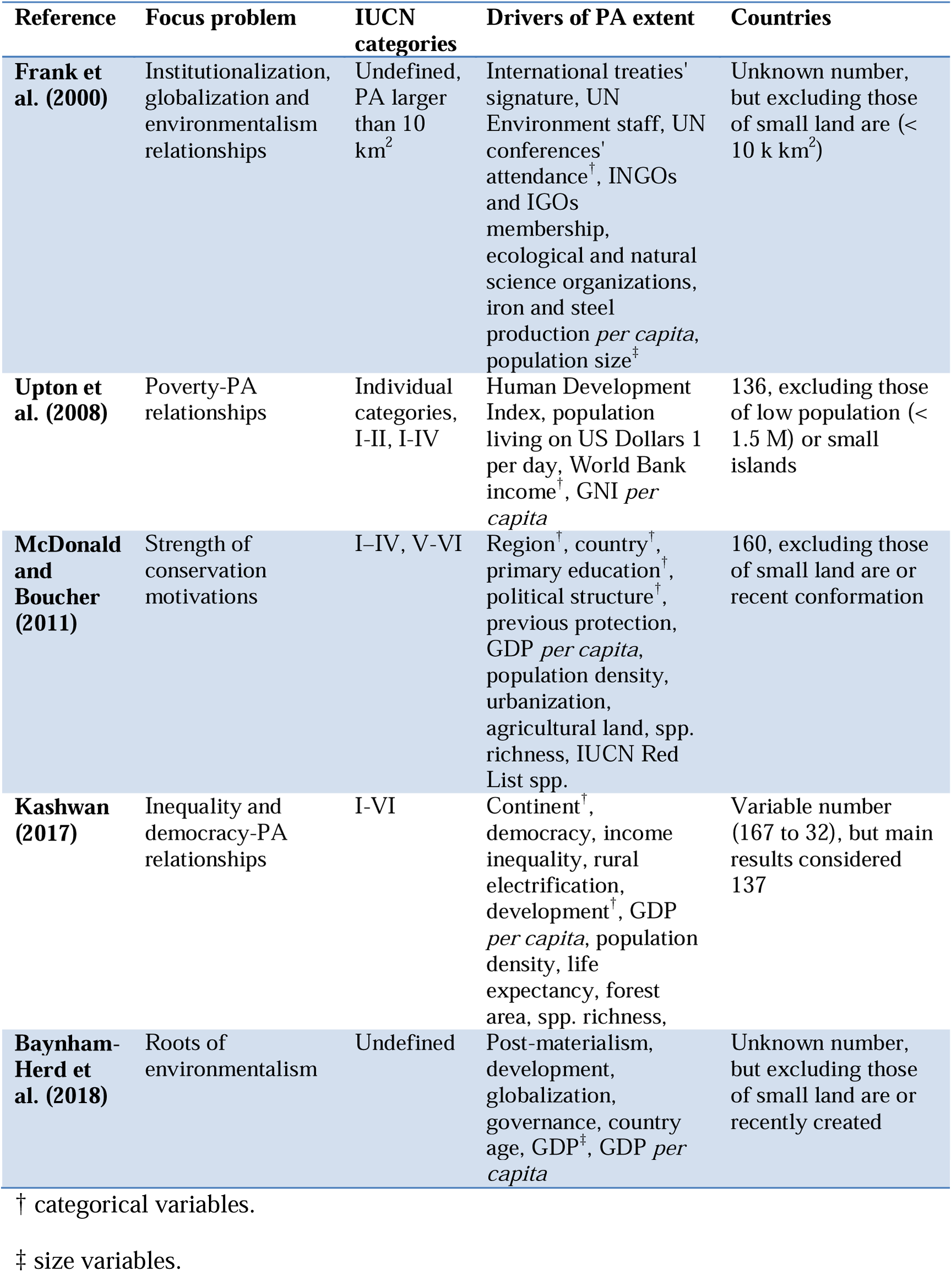
Non-exhaustive list of global studies about the cultural drivers of protected areas extent at a national level. Physical variables were excluded from the field of drivers (e.g., terrain slope). Acronyms: GDP Gross Domestic Product, GNI Gross National Income, IGOs intergovernmental organizations, INGOs international nongovernmental organizations, IUCN International Union for Conservation of Nature, PA protected areas, sd standard deviation, Spp. species, UN United Nations.

From the above studies, only Frank et al. (2000) and Baynham-Herd et al. (2018) marginally considered the size and power of countries in order to explain cross-national differences in nature conservation. Frank et al. (2000) included population size in models and Baynham-Herd et al. (2018) included economy size (Table 3). At this point, we identify a gap in knowledge. In a broad sense, country size and power (e.g., land area, economic activity, and military power; Crowards 2002b, Arvanitidis and Kollias 2016) constitute determining factors in numerous economic, political and social processes, such as market stability or social development (Alesina and Wacziarg 1998, Spolaore 2004, Prasad 2009). Given these circumstances, and returning to the initial question of the unbalanced distribution of assets in the world, the aim of this study is to explore the relationships between the intertwined concepts of size and power and the extent of protected areas. Additionally, we explore the relationship between the land area of a country and the size of the largest protected unit. Along with the size and power metrics, other cultural drivers used in previous research (e.g., education) were included in analyses in order to contextualize the shape and strength of relationships. Notably, the smallest and less powerful countries in the world are commonly excluded from comparative analyses in different fields, which biases our current knowledge to a partial spectrum of environmental or cultural conditions, therefore excluding extreme and deviant cases (Baldacchino and Milne 2006, Veenendaal and Corbett 2014). This study intends to amend these geographic, methodological and –perhaps– ideological biases by encompassing data from a more comprehensive set of countries.

## 2. Methods

The data about the size of individual protected areas (in km^2^) and of the extent of protected areas at a national level (in percentage) were obtained from the World Database on Protected Areas, March 2018 (IUCN and UNEP-WCMC 2018). We included only protected areas which have been specifically designated for nature protection, i.e., strict nature reserve, wilderness areas, national parks, natural monuments or features, and habitat/species management areas, categorized as I-IV under the IUCN guidelines (1994). In this database, protected areas in many countries (e.g., Bolivia, South Africa, Comoros) are almost exclusively labeled under the IUCN class “Not Reported”, which could lead to an underestimation of their national figures. In this sense, for these countries, we considered previous UNEP-WCMC categorizations (e.g., Bolivia or South Africa in 2013 had many protected areas categorized as I to IV) or included areas labeled with the general designations described above (e.g., national parks) and small variations of them. For those polygons that shared land and sea, we only considered the terrestrial area by subtracting the marine area to the overall GIS area from tabular data (Figure S1, Appendix 1). We excluded all protected areas with a “proposed” status. Due to a potential overestimation of national protected extent from overlapping problems (Deguignet et al. 2017), polygons were dissolved and new individual areas were recalculated using the Mollweide projection. Finally, for countries where polygonal data was partially or completely unavailable (> 50% of the units, e.g., Moldova), we included information from point data. After this data manipulation, the total global protected extent was 9.2 M km^2^, which is equivalent to 7.0% of the land surface, divided into approximately 114,000 units.

Samples of this study were the Parties of the Convention on Biological Diversity (CBD) (except for Monaco and the State of Palestine), including 191 countries members of the United Nations, Cook Islands and Niue (both under the Realm of New Zealand), and the United States of America (a United Nations member, but not a CBD party). Non-sovereign territories were excluded from analyses since many of the political decisions that concern their territorial management are held in central government administrations.

We related the protected extent at a national level to sixteen independent variables representing cultural drivers (Table 2). The first six variables are associated with the intertwined concepts of size and power of a country. The first variable is Crowards (2002b, a) classification based on non-hierarchical cluster analysis, generated from land area, gross domestic product (GDP) and population. The second, third and fourth variables are the individual land area, GDP and population. We added two more variables to the size and power group, i.e., the military expenditure and the possession of external territories (Baldacchino and Milne 2006, Arvanitidis and Kollias 2016). The following ten variables are related to general geographical, conservation and socio-economic characteristics of countries, and are used to contextualize the strength of variables related to size and power. These ten variables are equal or similar to those used in the global studies of Table 3, with the exception of the tourism contribution to GDP (variable #13) and whether the countries are continental or insular (variable #15). These last two variables are included as many local studies highlight the role of tourism as a driver of conservation (e.g., Maekawa et al. 2013), especially on islands (e.g., Sufrauj 2011). Finally, the governance value (variable #12) of Somalia was excluded by considering it as an outlier. All data is available in Appendix 2.

With an exclusively exploratory and descriptive purpose, we regressed protected extent (in percentage) to continuous variables by means of a Local Regression (LOESS) method. This non-parametric approach identifies patterns and fits a smoothed curve neither assuming any global function nor estimating a statistical significance of relationships (e.g., via a coefficient of determination) (Cleveland 1979). For categorical variables, we constructed violin plots, which are similar to box plots with a rotated kernel density plot on each side (Hintze and Nelson 1998). We also assessed the correlation between continuous variables through a Kendall’s τ non-parametric test (Whittaker 1987). The number of samples that were used for LOESS models (i.e., countries) varied according to data available (from 145 to 195, see Figure 2).

**Figure 1.**
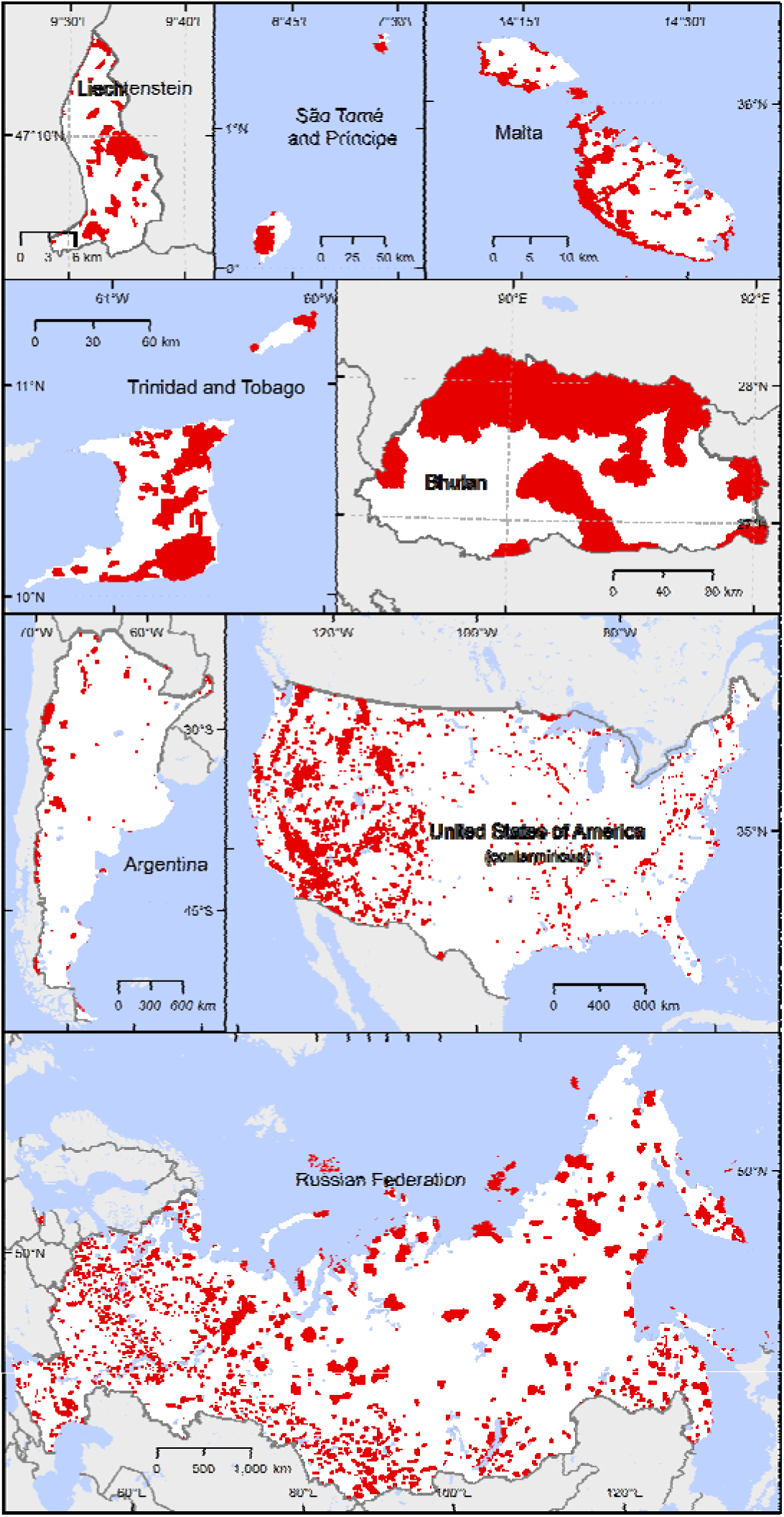
Protected areas categorized as I-IV under the International Union for Conservation of Nature guidelines (IUCN, 1994) in eight countries with contrasting geographies and human contexts. Alaska and Hawaii in the United States of America are not depicted.

**Figure 2.**
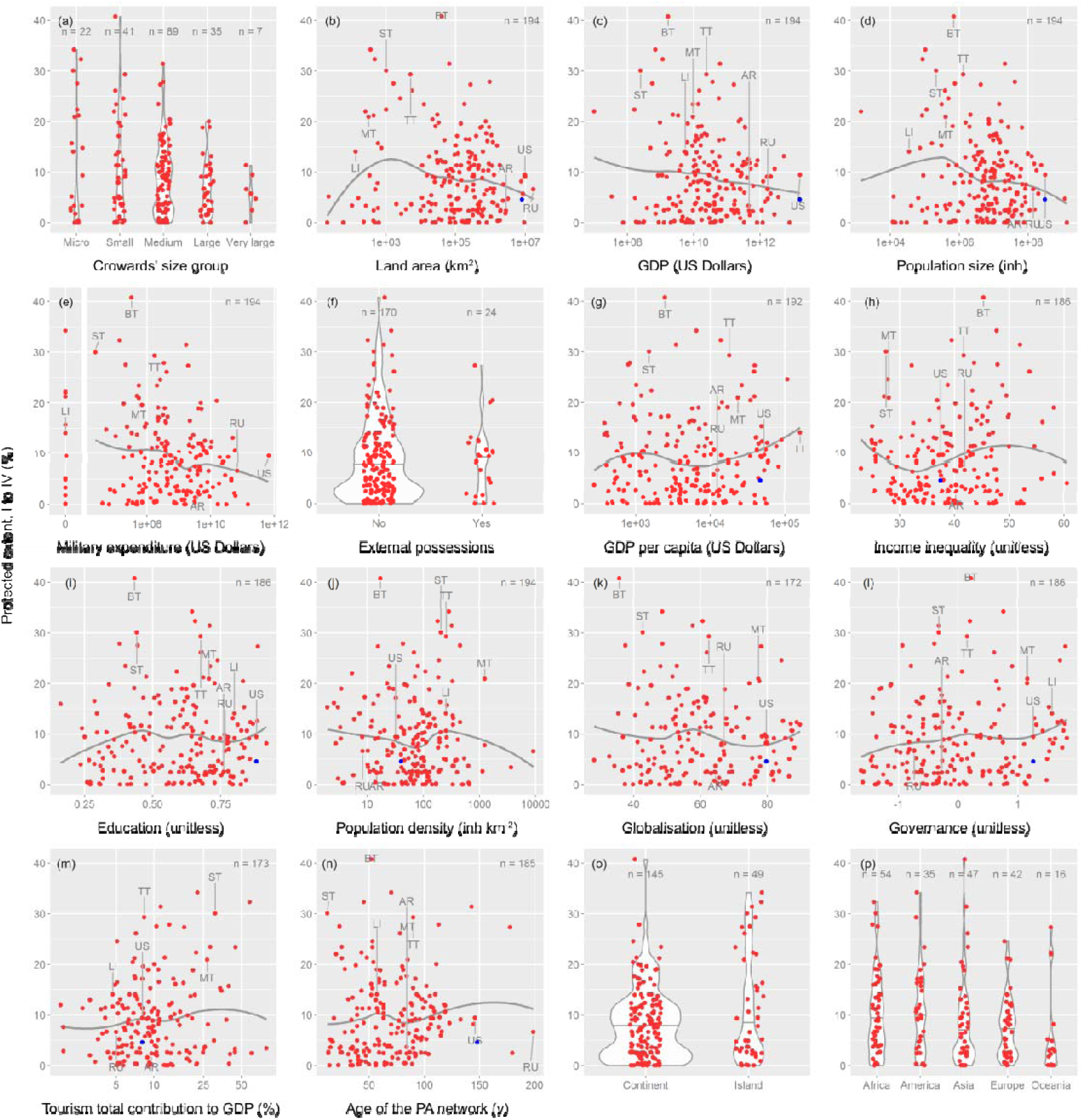
Extent and LOESS models of the networks of protected areas along socio-economical, conservation and geographical gradients. Panels (a) to (f) are related to the size and power of countries. In violin plots (a, f, o and p panels), horizontal lines represent the 0.5 quantiles. The number of samples is depicted on each panel. The eight countries of Figure 1 are labeled with the following acronyms: Argentina (AR), Bhutan (BT), Liechtenstein (LI), Malta (MT), Russia (RU), São Tomé and Príncipe (ST), Trinidad and Tobago (TT) and United States of America (US). Blue points represent the US conterminous states (these data do not feed LOESS models).

In order to measure the individual effect of the independent variables, we applied a random forest ensemble learning method (Breiman 2001), which estimates their importance (variable importance, VI) by looking at how much the mean square error (MSE) increases when the out-of-bag data (OOB) for that variable is permuted while all others are left unchanged (Liaw and Wiener 2002, Grömping 2009). To include all considered countries in the random forest, we filled missing values with the continental averages (5.4% of the values from the combination of 195 countries * 16 variables). Following Strobl et al. (2008, 2009) methodological suggestions, we calculated the VI using the “cforest” function of the “party” package (Hothorn et al. 2008). By selecting the subsample without replacement approach and the conditional permutation importance measure approach, VI measures can be used reliably even in situations where the independent variables are highly correlated and vary in their scale of measurement or their number of categories. In the cforest procedure, we chose a number of trees to grow, ntree, equal to 2500; a minimum size of the terminal nodes, nodesize, equal to 1; and a number of input variables at each split, mtry, equal to 3. For mtry, the chosen value minimized the OOB-MSE of the model. As VI results differed from run to run, we calculated a mean and standard deviation of VI values by running the model 50 times. The VI values were used here with an explanatory and interpretative rather than predictive aim. Data processing was conducted in RStudio v. 1.1.423 (RStudio Team 2018) (packages foreign, ggplot2, ggrepel, gridExtra and party).

## 3. Results

Land area is –perhaps– the most intuitive variable representing the size and power of a country. At a first cartographic glance, some of the largest countries of the world set aside a lower extent of their territories than some of the smallest ones for conservation (Figure 1). Russia and the United States of America, which are examples at the high end of the land area gradient, protect 6.6% and 9.4% of their territories, respectively. In contrast, Liechtenstein and São Tomé and Príncipe, which are examples at the low end of the land area gradient, protect 14.1% and 30.0% of their territories (Appendix 2). Figure 2 shows the protected extent along the classes or gradients of independent variables representing cultural drivers for all countries. Following Crowards’ classification (2002b, a), we show a high dispersion in the protected extent values within classes, with some “micro” to “large” countries achieving the highest values (up to 40.8% in Bhutan), but only one out of the seven “very large” countries surpassing 10% (Figure 2a). Complementing the cartographic description of Figure 1 regarding the land area, data reveal an inverse U-shaped curve pronounced at the low end of the gradient and with a gentle slope at the high end (Figure 2b). Very small countries protect little (e.g., Nauru, San Marino), but with a small increase in land area, some countries set aside significant portions of their territories to conservation (e.g., Liechtenstein, Niue). The maximum values of protected extent are found in small countries that exceed the ∼500 km^2^ (e.g., Luxembourg, Sri Lanka). Moving on along the land area gradient, the slope of the LOESS model becomes negative, as many countries have the potential to achieve high values (e.g., Tanzania, Chile), but many more have very low conservation values. Finally, the largest countries do not equate the values of the smallest: Out of the twenty most extensive countries, only two exceed a 10% protected (i.e., Indonesia and Mongolia).

The relationship between protected extent and other components of size and power also adopts an inverse U-shaped curve (i.e., population size) or negative linear shapes (i.e., military expenditure and gross domestic product –GDP) (Figure 2c-e). These consistent results would obey the strong and positive correlation that exists between the four continuous variables (Figure 3). More affluent countries protect less than less affluent ones: Out of the twenty countries with the largest GDP that account for more than 80.5% of global amounts, none achieve the 17% of protection suggested by the CBD convention. Conversely, out of the twenty countries with the lowest GDP, six does. Countries with higher military expenditure protect less than those with the lower ones: Out of the twenty countries with the largest military budget that accounts for more than 85% of global amounts, only Israel surpasses the 17% of protection. Conversely, out of the twenty countries with the smallest military expenditure, five surpass the 17% of protection. Albeit with a less clear pattern, those countries with external possessions set aside less land for conservation than those without these (Figure 2f).

**Figure 3.**
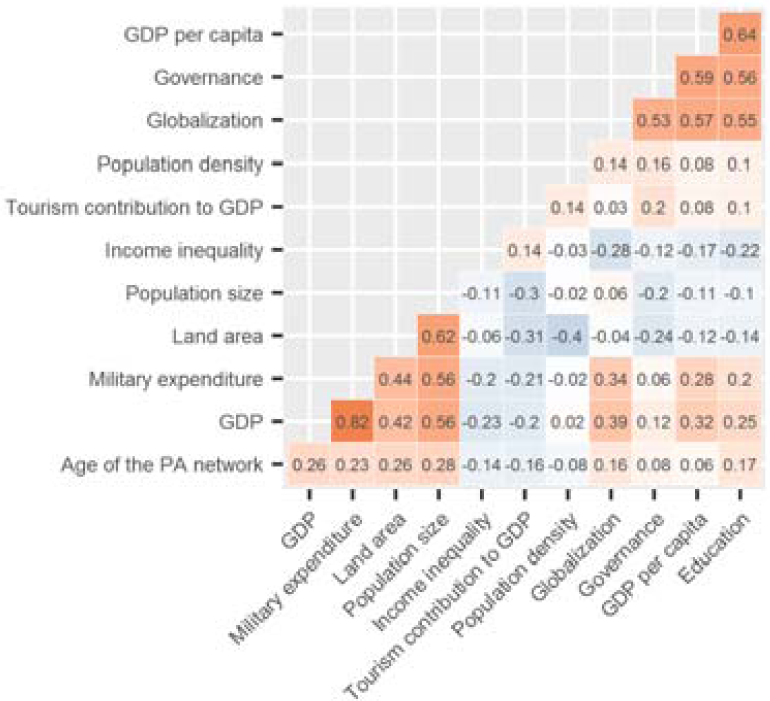
Kendall’s τ correlation coefficients between continuous variables (n = 194). Colours represent strength and sign (from positive red, to white, to negative blue).

According to the random forest analysis, the most relevant variable determining the protected extent of a country is income inequality, being ∼2.7 times more important than the following two variables, i.e., whether the country is continental or insular and military expenditure (VI = 1.37 ± 0.10, 0.54 ± 0.07 and 0.48 ± 0.06, respectively; Figure 4). The strong importance of income inequality obtains suggests the interaction with other variables, given the unclear dependence of the protected extent to this variable according to the LOESS model (Figure 2h). In comparison, other variables related or unrelated to size and power, such as the above mentioned land area (VI = 0.08 ± 0.06), governance (VI = 0.14 ± 0.05) or even the contribution of tourism to GDP (VI = −0.29 ± 0.06) suggest clearer relationships to protected extent from LOESS models (Figure 2b,l,m). Although none of the studies of Table 3 incorporated the condition of continental/insular among the driving factors, it is interesting to notice the high protected extent achieved by some insular countries (tropical, temperate or cold; Figure 2o). Out of the ten countries with the highest level of protection, eight are insular, and out of the next ten countries, half are insular.

**Figure 4.**
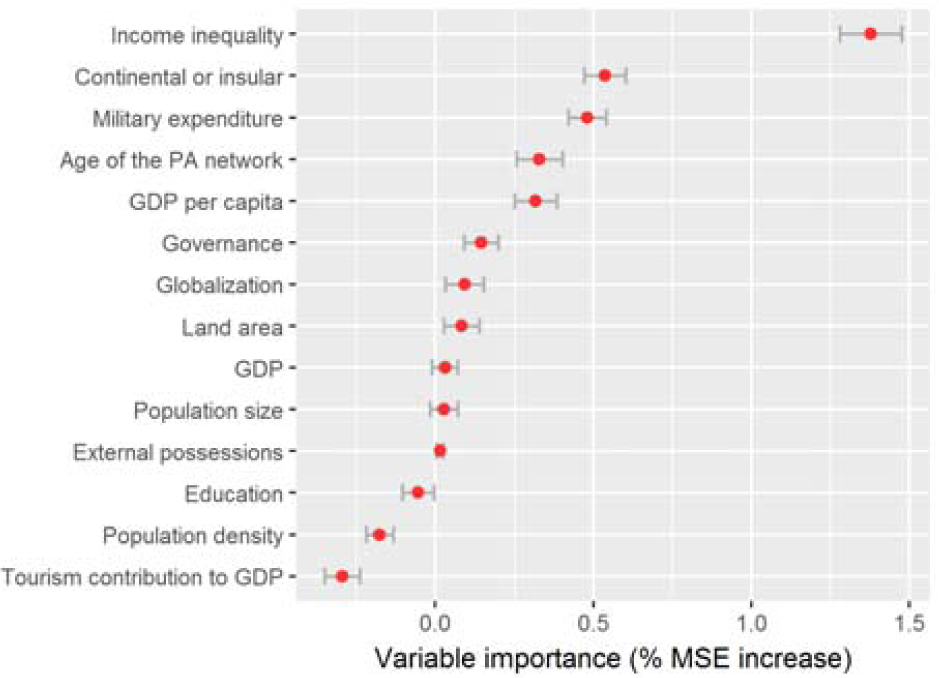
Relative importance of fourteen variables related to socio-economical, conservation and geographical contexts according to the random forest (RF). The variable importance is depicted by the increase in the mean square error when the out-of-bag data for a variable is permuted while all others are left unchanged. Variables #1 and #16 were excluded from the analysis.

In addition to an apparent effect of land area on the protected extent of a country, the smallest countries also preserve larger protected areas in relation to their particular size, with the Seychelles being at the top of the ranking (Figure 5). As other examples, we can mention the Ôbo Natural Park in the southernmost São Tomé island of São Tomé and Príncipe, which covers 24.4% of the 860 km^2^ of the country (Figure 1), while the Garsaelli Forest Reserve in Liechtenstein covers 6.7% of the national territory. Out of the first twenty countries whose most extensive protected area constitutes a larger fraction of the country, eight of them are “micro”, seven are “small” and five are “medium” following Crowards’ classification (2002b, a). The first “large” country that appears in the ranking is Venezuela, in the thirteenth position, closely followed by Chile (rank #34) and Algeria (rank #41).

**Figure 5.**
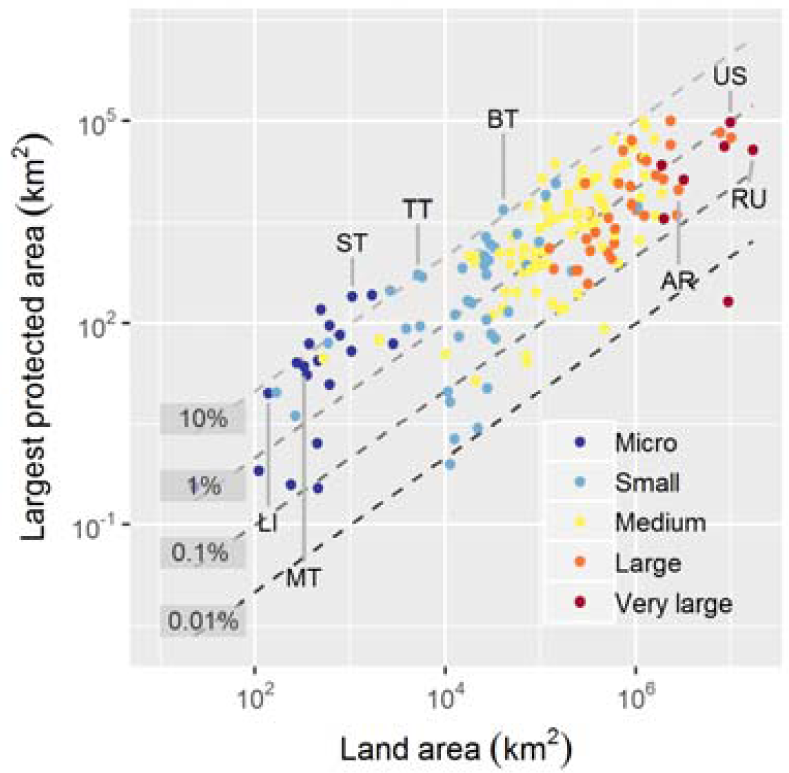
Relationship between the largest protected area of the country and its land area. The four lineal models represent isolines of this relationship (e.g., the 10% isoline represents a single protected area that occupies a tenth of the land area). Different colors represent the size classes of Crowards (2002b, a) which consider, besides land area, population and GDP. See country acronyms in Figure 2.

## 4. Discussion

There is a broad consensus in society that protected areas are one of the most effective tools in the conservation of nature, and that it is necessary to extend their current surface and connect all biophysical systems (Rodrigues et al. 2004, Watson et al. 2014, Saura et al. 2018). However, this globalized discourse contrasts with the results of this study for the <u>more developed or powerful</u> countries, as it is argued in section 4.1. A key to forecast scenarios and eventually balance efforts across countries lies in the comprehension of the cultural drivers of conservation, as it is argued in section 4.2.

### 4.1. Size and power

The LOESS model in Figure 2b suggests that the larger and more powerful countries (in terms of land area) maintain a conservation debt regarding protected extent in comparison with micro-to medium-size countries. Large to very large countries are also mostly affluent, populated and powerful (Figure 3), a conjunction of factors that make them fundamentally responsible for the conservation of nature. As a matter of fact, these countries manage a remarkable amount of natural resources and biodiversity, occupy different continents or hemispheres, possess the material resources to maintain and promote conservation programs, and shape and determine their own and others’ economic and political actions with the greatest independence (Neumann and Gstöhl 2004, Beckley 2018). Certainly, every country should make a similar attempt to conserve a fraction of their natural resources (SCBD 2010), as the fulfillment of common goals by small countries ensures the protection of their geographical or biological singularities. However, this legal equality has a political and –essentially– ecological counterpart, since an effective growth of the global protected area network depends on larger and more powerful countries which have the actual capacity to generate radical changes at that spatial level. Jenkins and Joppa (2009) stated that the global increase in protected extent during the 2000 decade was accounted for mostly by Brazil’s protected area expansion.

More than two decades ago, Wells and Williams (1998) pointed out that, with the demise of communism in Russia, the economic resources allocated to conservation sharply declined, with a consequent weakening of law enforcement and an increase in illegal activities in protected areas. Confirming and extending these findings, Watson et al. (2014) suggested that there was significant evidence that more affluent and extensive countries such as Australia, the United States of America or Canada were cutting financial and human resources for the conservation sector, and were even overlooking existing conservation policies and legislation. This has occurred in spite of the strong discourse in these countries towards increasing the size and effectiveness of protected area networks. Several studies endorse our findings by considering the level of anthropization of protected areas (Leroux et al. 2010, Jones et al. 2018), the specific efforts in the conservation of a taxon (Lindsey et al. 2017), or even the attitudes of the population (Nawrotzki 2012). By means of social surveys, Nawrotzki (2012) stated in fact that the strongest opposition toward environmental protection was observed in conservative people of more powerful, capitalist countries, while in peripheral, less developed ones, conservative people were more environmentally friendly than liberal ones.

This differential effort emerges so markedly that it is interesting to return to a selective and meaningful comparison between extreme cases. If Russia and the United States of America sought to repeat the examples of Liechtenstein and São Tomé and Príncipe, respectively, their protected area network would need to incorporate 1.4 M km^2^ and 2.0 M km^2^, respectively, to their current networks. In this speculative exercise, the inclusion of these 3.4 M km^2^ would increase the global protected extent by 2.6%, from 7.0% to 9.6% under I-IV categories under IUCN guidelines (1994). If we extend comparisons to the size of the largest protected area (Figure 5), Russia’s largest protected area would need to have 30.8 times its current size, while the United States of America’s largest protected area would need to have 25.3 times its current size. This examination partially supports and extends what was suggested by Upton et al. (2008), i.e., that wealthy countries have protected areas of smaller size than poorer countries. Comparisons can be considered at some point unlikely due to the deep economic and social implications that Russia and the United States of America (as well as other countries like China or Indonesia) would have to face in light of a different territorial order. However, comparisons stress that some small to medium-sized countries have decided to follow conservation-prone spatial planning without many –at least financial– apparent difficulties. Renowned Bhutan’s conservation efforts exceed protected areas, as the constitution mandates that at least 60% of the country must remain with its natural forest cover, while the government has committed to remaining carbon neutral (Wangchuk 2007, Lham et al. 2019).

Turning back to the Aichi agreements, the debt held by the largest or most powerful countries is enlarged by considering the second clause of Target 11 (SCBD 2010), which sets that protected networks have to sample all natural conditions and all levels of life organization with the same effort. According to Barr et al. (2011), the very large countries from Crowards’ classification (2002b, a) protect in a divergent way: Russia’s network equitably encompasses most ecoregions of the country, occupying position #12 of 83 in the ranking, while Brazil, the United States of America, Indonesia, India, Mexico, and China surpass position #55. To reinforce this idea, we deliberately excluded the non-contiguous states of Alaska and Hawaii of the United States of America, as well as their external possessions in Figure 1. Conterminous United States of America reaches a protected extent of 4.6% under the I to IV categories, a value lower than those of 121 countries. These biases can be attributed to the fact that protected areas are frequently established on territories that face little human interventions and have comparatively low opportunity-costs, at least at the time of their establishment (Joppa and Pfaff 2009, Baldi et al. 2017, Baldi et al. 2019). One possibility to overcome these problems of imbalances among and within countries is that international representation goals are not established at the country level but at the level of ecoregion or physical environments (Aksenov et al. 2015, Baldi et al. 2019). In this way, larger and more powerful and of smaller and less powerful countries should agree on multilateral strategies of conservation of their natural conditions regardless of the protected national extent.

### 4.2. Size and power in context

The main lesson from Figure 2 is that none of the individual relationships explored was particularly strong. This fact does not detract what was expressed in section 4.1 but rather indicates that there are multiple interactions among variables not identified in the analyses. Interactions would explain why income inequality has prevailed in the random forest ranking (Figure 4). In fact, Kashwan (2017) found a strong interaction between income inequality and the system of government: When democracy prevailed, inequality led to less protection, whereas when totalitarianism did, inequality led to more protection.

Including income inequality and other variables in these analyses was aimed at contextualizing the importance of size and power drivers. From these, the one that reached the greatest importance turned out to be military expenditure (third position), followed by land area (eighth position, with values similar to globalization and governance), and GDP, population size and external possessions in ninth, tenth and eleventh positions, respectively (Figure 4). Regardless of the position in the ranking, the question about which direct or indirect mechanisms explain the effect of size and power remain unanswered. Having stated this, the empirical knowledge generated in this and previous studies (see Table 3) provides evidence to propose the following *a posteriori* hypotheses:

(#1) The *per capita* costs of different public goods and services are determined by the number of taxpayers of a country (Alesina 2003, Spolaore 2004), as well as by other elements of tax structure. In this regard, the larger and more powerful the country is (in population terms), the more economically viable it would be to maintain and expand a network of protected areas.

(#2) Integration to world society is commonly related to economic wealth in smaller or less powerful countries (Pelling and Uitto 2001, Prasad 2009). In this regard, and as opposed to the previous hypothesis, conservation would be internally boosted by accomplishing multilateral or international treaties, regulations, or agreements (like the CBD, Woodley et al. 2012), or by attending transnational social movements (Lewis 2000). In addition to this integration, conservation projects in smaller countries frequently obtain financial assistance from international non-governmental organizations (NGOs) (Frank et al. 2000, Kashwan 2017). This can eventually be associated with a green grabbing process (Fairhead et al. 2012) or a sign of soft power (i.e., the ability to attract and co-opt others) that larger countries exert over smaller ones (Arvanitidis and Kollias 2016, Beckley 2018).

(#3) The diversity and abundance of natural resources are generally determined by the land area of a country through a sampling effect (except those occupying extreme deserts) (Freudenberger et al. 2012). In this regard, the larger the country is (in terms of land area), the greater the redundancy of diverse natural resources will be. Given this redundancy, larger countries would allocate a small fraction of their territories to conservation in order to achieve the representation goals. Complementarily, smaller countries would allocate a large fraction of their territories in order to maintain their natural system, ensuring the provision of varied resources or services and the achievement of representation goals.

(#4) Given the interaction between the aspects associated with an economy of scale (hypothesis #1) and a limited diversity of resources (hypothesis #3), the economic viability of smaller and less powerful countries (in terms of land area, population, GDP) would be conditioned by non-extractive or unconventional industries, such as tourism in tropical islands (Croes 2013). In this regard, the smaller and less powerful is the country, the tourism industry would be boosted by the creation of infrastructure and the maintenance of landscape quality and biological diversity in protected areas.

Considering the results depicted in Figures 1 and 2a-f, the mechanism that supports the first of the four hypotheses would not prevail, since even countries with a very low population maintain extensive protected area systems (e.g., Iceland, Mongolia, Namibia) (Figure 2d,j). The other three hypotheses could advocate mechanisms that have effective implications for conservation, although these have received mixed support from our results. As an example, we found that the highly correlated governance-globalization drivers (Figure 3), which achieved an intermediate position in the importance ranking (Figure 4), showed an inconclusive relationship with the protected extent (Figure 2k,l). This questions the interactions or mediations of hypothesis #2. Results also question the findings of Frank et al. (2000) and Baynham-Herd et al. (2018), who emphasized the positive role of social and cultural processes that promote the links to world society (i.e., globalization) directing conservation efforts. As another example considering hypothesis #4, countries of small to intermediate size and economically dependent on tourism activities can have extensive protected area systems (e.g., Cambodia, Malta), or cannot (e.g., Georgia, Maldives).

Other less explored factors, such as the closeness to natural environments, the social sense of belonging, cohesion or self-sufficiency, as well as the degradation, scarcity or finitude of natural resources in resource-constrained environments, could support the thesis of a greater conservation effort in smaller or less powerful countries (Aguilera-Klink et al. 2000, McNeely 2015). Oceanic islands could be used to test these ideas, as they frequently maintain well-organized strategies of land management due to their high environmental, demographic, and economic vulnerabilities (Pelling and Uitto 2001, Christensen and Mertz 2010). We find that the condition of insularity (second in the importance ranking) could support these postulates since violin plots in Figure 2o suggest that protection in insular countries is larger than that in continental countries.

Our results could be used to unfold conservation scenarios in the face of geopolitical changes, specifically in relation to the number and size of countries. Two opposite phenomena occur in this regard. On the one hand, new countries will probably emerge as there are still subnational regions and stateless nations asserting for full sovereignty and recognition (e.g., New Caledonia, Kurdistan) (Baldacchino and Hepburn 2012, Veenendaal and Corbett 2014). On the other hand, the pursuit of sovereignty has sharply declined in the last decades (e.g., Basque Country, Scotland) (Baldacchino and Milne 2006, Baldacchino and Hepburn 2012), some once sovereign and recognized countries have been (re)united (e.g., Germany, Yemen) and political blocks have been formed with different levels of supranational unification (e.g., Turkic Council, European Union). If small territories gained sovereignty, we wonder whether the creation of protected areas would slow down (hypothesis #1) or accelerate (hypotheses #2 to #4). And *vice versa*, if federations, confederations, and other unions consolidate, it would be interesting to question whether the creation of protected areas would accelerate (hypothesis #1) or slow down (hypotheses #2 to #4).

Finally, we should highlight some caveats of the paper. First, the land area could be intuitively considered a geographical –more than a cultural– trait. However, as stated by Alesina (2003), the land area would not be necessarily a factor external to a country’s culture, as the same culture could determine this merely geographical feature. Second, we have exclusively evaluated the relationships of each current <u>cultural driver</u> and the protected extent. However, in the historical process of growth of protected area networks, these relationships have varied (McDonald and Boucher 2011, Radeloff et al. 2013). At last, the role of unconsidered protected areas could modify the messages from this study. By comparing previous values presented in this text, protected areas under V, VI or miscellaneous IUCN categories encompass approximately 126,000 units and 5.7% of the global protected area. Nevertheless, we left aside these categories (and other such as wilderness areas, Dietz et al. 2015) as their governance and tenure potentially lack formal protection and management, they have uncertain conservation objectives and long-term capabilities, and their enforcement of law is compromised (Shafer 2015, 2019).

## 5. Conclusions

The commitment of countries to nature conservation, specifically through the deployment of protected areas, showed to be greatly uneven among countries, from those which completely devoid of this legal figure to those in which nearly half of the territory is strictly protected. These differences would obey to the interaction of several underlying and direct cultural drivers. Previous studies that observe these relationships excluded countries of small size, sparsely populated, recently conformed or of insular character, omitting thus extreme and deviant geographical or cultural cases. Meanwhile, until now the size and power of a country as a driver of the protected extent of a country had not been explored. We intended to amend both situations, finding that the largest or most powerful countries have made a lower conservation effort than the smaller or less powerful ones. Size and power would mediate individual or joint effect of other drivers or would act directly as a mechanism by allowing, for example, stronger countries to impose policies over weaker countries or by abstaining to participate (the stronger) countries in international agreements.

Regardless of these points, the largest and most powerful countries are the ones that have the greatest responsibility in nature protection, given that internal changes in their conservation policies aimed at increasing the extension and financing their protected areas imply the success of the global conservation of nature. Perhaps a more plausible conclusion about the differences among countries is that the larger and more powerful ones protect less, not because they cannot do better, but because it is not part of their political agendas. Paraphrasing Lewis and Wigen (1997), an increasingly integrated world demands a more modest, honest, and accurate geographical depiction of the distribution of protected areas to understand the needs, debts, and opportunities in the conservation of nature.

## Acknowledgments

I would like to thank S.A. Schauman, S. Aguiar, J.I. Whitworth-Hulse, O.A. Martín, T. Milani, E.G. Jobbágy and M. Grainger for their ideas and collaboration in different stages of the study. We would also like to thank D. McGreevey and the Gabinete de Asesoramiento en Escritura Científica en Inglés, UNSL, for her service and help.

