## Appendix_1 for "Nature protection across countries: Do size and power matter?"

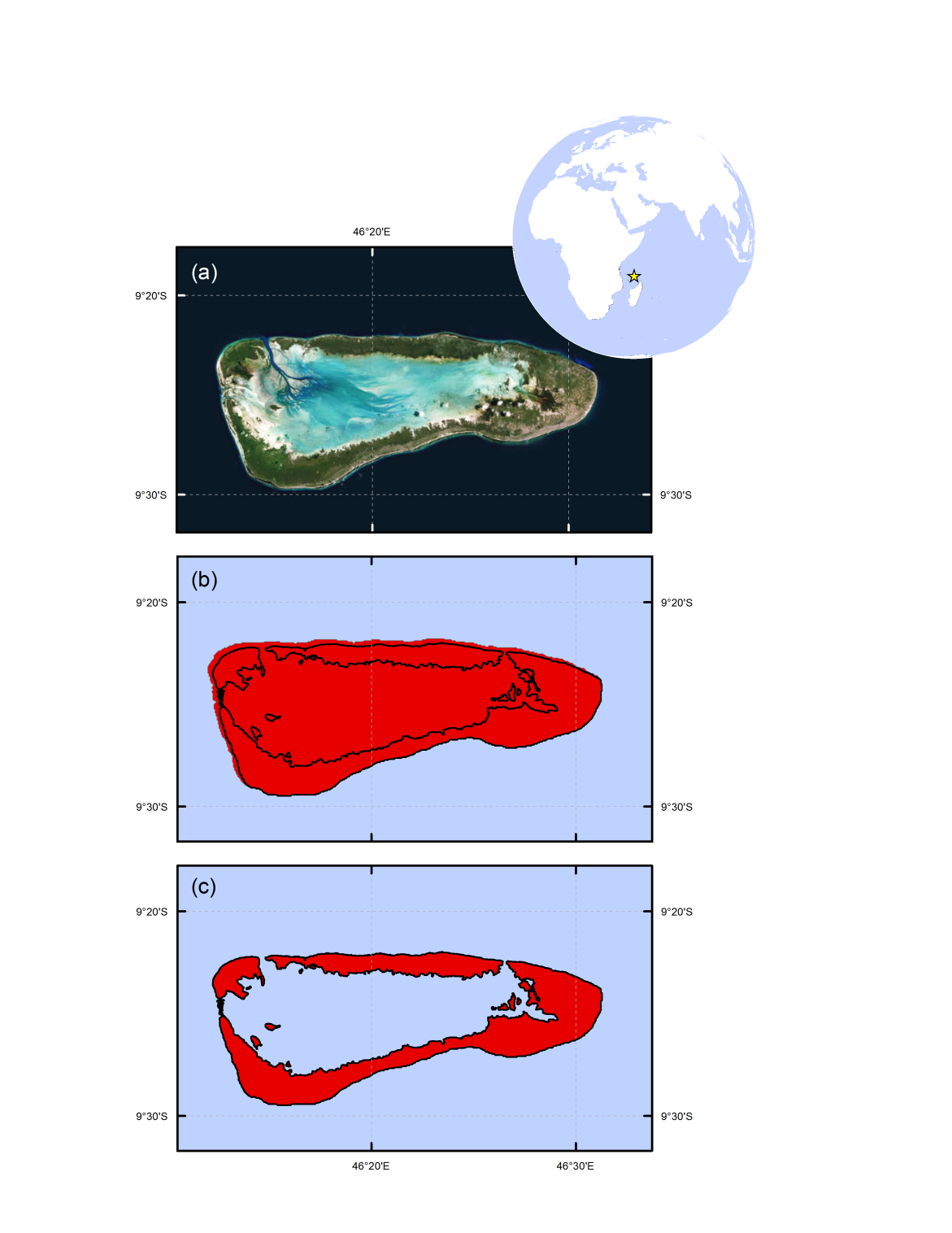

Figure S1. Example of data manipulation in mixed land/marine protected areas. In (a) a high spatial resolution image from Google Earth of Aldabra atoll, Seychelles. The ring-shaped island encloses a lagoon of shallow sea water (light blue). In (b) the protected area (in this case, Special Reserve) according to the IUCN and UNEP-WCMC database (2018). In (c) A graphical exercise of the subtraction of marine area to the overall area.

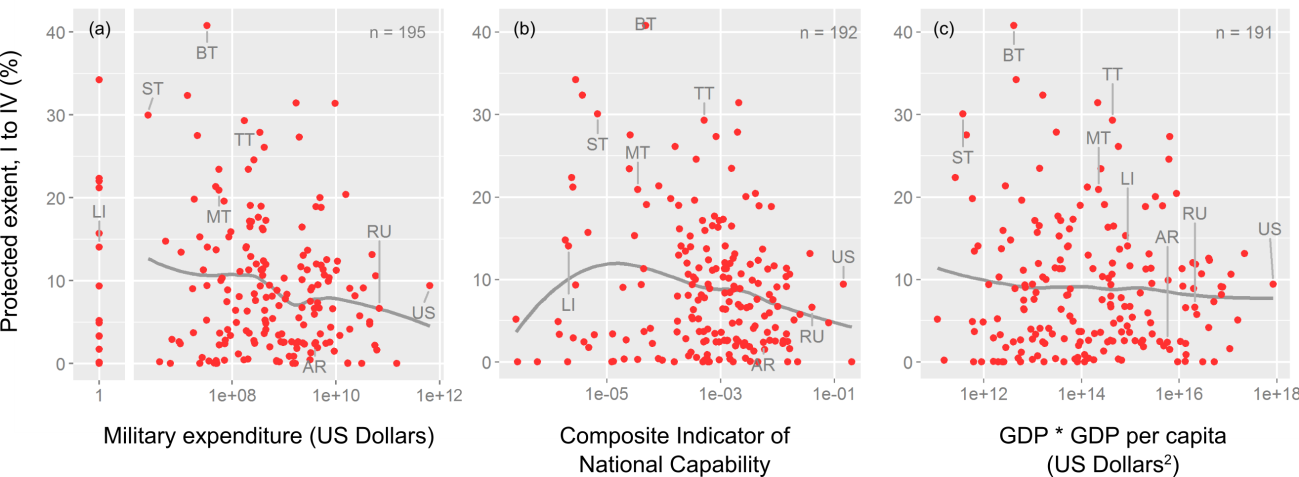

Figure S2. Extent and LOESS models of the networks of protected areas along national power gradients. In (a) military expenditure, in (b) the Composite Indicator of National Material Capabilities v. 5.0 (Singer 1988) and in (c) the interaction among GDP and GDP per capita (Beckley 2018). The number of samples is depicted on each panel. The eight countries of Figure 1 are labeled with the following acronyms: Argentina (AR), Bhutan (BT), Liechtenstein (LI), Malta (MT), Russia (RU), São Tomé and Príncipe (ST), Trinidad and Tobago (TT) and United States of America (US).
