## Appendix_2 for "Nature protection across countries: Do size and power matter?"

| Country | Protected extent  (%) | Size of largest protected area (h km^2^) | Crowards' country size | Land area  (k km^2^) | Gross domestic product (M US Dollars) | Population size  (M inh) | Military expenditure  (M US Dollars) | External possessions | GDP *per capita*  (US Dollars * inh^-1^) | Income inequality (unitless) | Education  (unitless) | Population density  (inh * km^-2^) | Globalization  (unitless) | Governance  (unitless) | Tourism contribution to GDP (%) | Age of the PA network | Continental or insular | Continent |
| --- | --- | --- | --- | --- | --- | --- | --- | --- | --- | --- | --- | --- | --- | --- | --- | --- | --- | --- |
| Afghanistan | 2.75 | 8.08 | Me | 642.18 | 16649 | 28.4 | 261.1 | N | 600 | 33.9 | 0.32 | 44.2 | 36.7 | -1.63 | NA | 45 | C | As |
| Albania | 13.93 | 8.71 | Sm | 28.34 | 12197 | 3.6 | 183.6 | N | 4262 | 38.7 | 0.65 | 128.4 | 63.8 | -0.16 | 21.3 | 52 | C | Eu |
| Algeria | 4.35 | 993.31 | La | 2308.86 | 176000 | 34.2 | 7962.8 | N | 4925 | 29.2 | 0.58 | 14.8 | 53.6 | -0.85 | 6.7 | 46 | C | Af |
| Andorra | 21.18 | 0.28 | Mi | 0.45 | 3395 | 0.1 | 0.0 | N | 39310 | 27.2 | 0.69 | 185.5 | NA | NA | 7.0 | 25 | C | Eu |
| Angola | 6.64 | 153.41 | Me | 1244.65 | 97209 | 12.8 | 3849.1 | N | 4135 | 43.7 | 0.43 | 10.3 | 42.7 | -1.04 | 4.2 | 80 | C | Af |
| Antigua and Barbuda | 2.39 | 0.00 | Mi | 0.45 | 1271 | 0.1 | 10.0 | N | 12841 | 47.9 | 0.69 | 189.7 | NA | 0.71 | 60.4 | 13 | I | Am |
| Argentina | 2.37 | 93.79 | La | 2784.31 | 469000 | 40.9 | 3912.6 | N | 12443 | 40.4 | 0.76 | 14.7 | 64.5 | -0.27 | 9.6 | 84 | C | Am |
| Armenia | 12.37 | 15.12 | Sm | 29.59 | 10339 | 3.0 | 398.7 | N | 3643 | 34.3 | 0.70 | 100.3 | 61.5 | -0.27 | 14.0 | 60 | C | As |
| Australia | 8.18 | 669.66 | La | 7691.17 | 1250000 | 21.3 | 23274.6 | Y | 59755 | 33.1 | 0.92 | 2.8 | 79.6 | 1.59 | 10.9 | 145 | C | Oc |
| Austria | 8.88 | 8.05 | Me | 83.99 | 410000 | 8.2 | 3258.0 | N | 48312 | 27.9 | 0.77 | 97.7 | 87.4 | 1.53 | 15.6 | 77 | C | Eu |
| Azerbaijan | 10.12 | 12.03 | Me | 86.25 | 55506 | 8.2 | 2217.4 | N | 6525 | 28.5 | 0.68 | 95.5 | 56.5 | -0.78 | 14.6 | 89 | C | As |
| Bahamas | 23.41 | 1.35 | Sm | 12.59 | 9493 | 0.3 | 55.0 | N | 26683 | 48.8 | 0.71 | 24.6 | 48.0 | 0.91 | 44.8 | 53 | I | Am |
| Bahrain | 7.78 | 0.51 | Sm | 0.58 | 28485 | 0.7 | 1051.9 | N | 23125 | NA | 0.68 | 1244.2 | 66.2 | -0.01 | 9.9 | 39 | I | As |
| Bangladesh | 2.60 | 6.60 | La | 136.90 | 139000 | 156.1 | 1868.7 | N | 1008 | 38.8 | 0.40 | 1139.9 | 44.3 | -0.86 | 4.3 | 38 | C | As |
| Barbados | 0.35 | 0.02 | Mi | 0.44 | 4568 | 0.3 | 49.2 | N | 16245 | 47.8 | 0.73 | 640.7 | 58.5 | 1.12 | 39.9 | 56 | I | Am |
| Belarus | 6.96 | 21.59 | Me | 207.50 | 59812 | 9.6 | 802.0 | N | 6675 | 23.0 | 0.78 | 46.5 | 57.8 | -0.81 | 5.9 | 93 | C | Eu |
| Belgium | 11.84 | 1.31 | Me | 30.67 | 496000 | 10.4 | 5165.0 | N | 44628 | 25.6 | 0.81 | 339.6 | 89.9 | 1.31 | 5.6 | 119 | C | Eu |
| Belize | 9.04 | 10.73 | Sm | 22.30 | 1526 | 0.3 | 17.2 | N | 4667 | 46.7 | 0.68 | 13.8 | 53.8 | -0.15 | 38.1 | 34 | C | Am |
| Benin | 7.46 | 58.19 | Me | 116.11 | 7887 | 8.8 | 78.9 | N | 836 | 41.1 | 0.34 | 75.7 | 48.4 | -0.27 | 5.8 | 76 | C | Af |
| Bhutan | 40.77 | 49.40 | Sm | 40.37 | 1696 | 0.7 | 32.5 | N | 2472 | 45.3 | 0.43 | 17.1 | 35.7 | 0.21 | NA | 52 | C | As |
| Bolivia | 17.27 | 17.28 | Me | 1086.81 | 24829 | 9.8 | 446.8 | N | 2751 | 45.1 | 0.63 | 9.0 | 61.1 | -0.58 | 7.1 | 79 | C | Am |
| Bosnia and Herzegovina | 0.55 | 1.61 | Me | 40.03 | 17541 | 4.6 | 201.4 | N | 4862 | 40.1 | 0.66 | 115.2 | 67.9 | -0.32 | 9.2 | 64 | C | Eu |
| Botswana | 18.98 | 524.76 | Me | 579.03 | 13649 | 2.0 | 361.4 | N | 6997 | 58.1 | 0.61 | 3.4 | 56.4 | 0.68 | 10.9 | 57 | C | Af |
| Brazil | 5.74 | 423.21 | VL | 8472.66 | 2060000 | 198.7 | 29054.9 | N | 11191 | 46.0 | 0.61 | 23.5 | 60.6 | 0.00 | 8.5 | 92 | C | Am |
| Brunei | 26.09 | 4.99 | Sm | 5.72 | 14818 | 0.4 | 412.6 | N | 39139 | 53.7 | 0.68 | 67.9 | 61.9 | 0.60 | 7.2 | 78 | I | As |
| Bulgaria | 2.61 | 7.78 | Me | 112.76 | 52892 | 7.2 | 878.0 | N | 7432 | 33.0 | 0.73 | 63.9 | 76.7 | 0.19 | 12.8 | 87 | C | Eu |
| Burkina Faso | 11.12 | 182.61 | Me | 272.77 | 10082 | 15.7 | 141.4 | N | 646 | 35.3 | 0.24 | 57.7 | 49.5 | -0.37 | 3.8 | 82 | C | Af |
| Burundi | 3.66 | 5.37 | Sm | 27.04 | 2344 | 9.0 | 57.4 | N | 277 | 33.6 | 0.31 | 332.4 | 34.2 | -1.17 | 5.5 | 84 | C | Af |
| Cambodia | 23.44 | 40.95 | Me | 181.06 | 13780 | 14.5 | 203.4 | N | 1025 | 38.8 | 0.40 | 80.1 | 52.9 | -0.79 | 28.3 | 25 | C | As |
| Cameroon | 11.30 | 53.00 | Me | 464.32 | 28980 | 18.9 | 360.1 | N | 1416 | 39.6 | 0.43 | 40.7 | 48.1 | -0.93 | 8.1 | 50 | C | Af |
| Canada | 9.07 | 559.82 | La | 9945.53 | 1630000 | 33.5 | 18473.6 | N | 48622 | 31.4 | 0.85 | 3.4 | NA | 1.63 | 6.3 | 133 | C | Am |
| Cabo Verde | 2.64 | 0.85 | Sm | 3.88 | 1720 | 0.4 | 9.4 | N | 3347 | 43.1 | 0.50 | 110.6 | 49.8 | 0.53 | 44.5 | 13 | I | Af |
| Central African Republic | 14.04 | 190.92 | Me | 617.98 | 1861 | 4.5 | 33.2 | N | 410 | 47.1 | 0.30 | 7.3 | 35.4 | -1.43 | 6.4 | 93 | C | Af |
| Chad | 9.42 | 835.12 | Me | 1266.28 | 11079 | 10.3 | 522.3 | N | 902 | 37.9 | 0.25 | 8.2 | 36.8 | -1.37 | 4.1 | 63 | C | Af |
| Chile | 20.01 | 358.21 | La | 736.60 | 229000 | 16.6 | 4911.4 | Y | 14458 | 46.6 | 0.73 | 22.5 | 74.7 | 1.15 | 10.1 | 111 | C | Am |
| China | 0.01 | 2.11 | VL | 9374.57 | 7780000 | 1338.6 | 148000.0 | Y | 6784 | 51.6 | 0.55 | 142.8 | 59.9 | -0.52 | 9.0 | 33 | C | As |
| Colombia | 11.25 | 279.85 | La | 1135.12 | 301000 | 45.6 | 10127.4 | N | 7022 | 50.0 | 0.57 | 40.2 | 59.9 | -0.31 | 5.8 | 44 | C | Am |
| Comoros | 27.51 | 2.68 | Mi | 1.67 | 565 | 0.8 | 21.4 | N | 797 | 47.9 | 0.45 | 450.0 | 37.5 | -0.93 | 10.5 | 23 | I | Af |
| Cook Islands | 0.17 | 0.00 | Mi | 0.24 | 183 | 0.0 | 0.0 | N | NA | NA | NA | 50.3 | NA | NA | NA | 40 | I | Oc |
| Costa Rica | 11.99 | 19.96 | Me | 51.14 | 42660 | 4.3 | 293.0 | N | 10260 | 45.3 | 0.63 | 83.2 | 67.8 | 0.61 | 13.4 | 63 | C | Am |
| Côte d'Ivoire | 6.39 | 117.56 | Me | 320.68 | 28232 | 20.6 | 425.1 | N | 1380 | 37.4 | 0.35 | 64.3 | 53.1 | -0.97 | 7.5 | 86 | C | Af |
| Croatia | 2.54 | 2.98 | Me | 55.08 | 58615 | 4.5 | 972.9 | N | 13160 | 27.9 | 0.71 | 81.5 | 77.1 | 0.42 | 24.7 | 25 | C | Eu |
| Cuba | 4.67 | 52.06 | Me | 109.93 | 70321 | 11.5 | 2376.5 | N | 6598 | 38.0 | 0.74 | 104.2 | 57.1 | -0.57 | 9.6 | 17 | I | Am |
| Cyprus | 3.61 | 0.92 | Sm | 5.40 | 24306 | 0.5 | 412.1 | N | 27665 | 30.4 | 0.73 | 98.5 | 78.0 | 1.04 | 21.4 | 35 | I | As |
| Czech Republic | 2.47 | 6.85 | Me | 78.76 | 207000 | 10.2 | 2313.1 | N | 19589 | 25.5 | 0.83 | 129.7 | 81.7 | 0.92 | 7.8 | 180 | C | Eu |
| Democratic Republic of the Congo | 7.57 | 436.85 | La | 2325.24 | 26756 | 68.7 | 281.1 | N | 405 | 38.5 | 0.39 | 29.5 | NA | NA | 1.9 | 113 | C | Af |
| Denmark | 12.01 | 2.92 | Me | 42.70 | 329000 | 5.5 | 4185.6 | Y | 58372 | 25.0 | 0.87 | 128.8 | 87.1 | 1.79 | 7.7 | 118 | C | Eu |
| Djibouti | 0.01 | 0.03 | Sm | 21.85 | 1265 | 0.5 | 48.7 | N | 1586 | 38.4 | 0.27 | 23.6 | 52.9 | -0.71 | NA | 14 | C | Af |
| Dominica | 14.76 | 0.67 | Mi | 0.77 | 499 | 0.1 | 5.2 | N | 7142 | 43.0 | 0.61 | 94.9 | 52.8 | 0.69 | 34.7 | 43 | I | Am |
| Dominican Republic | 16.30 | 11.56 | Me | 48.44 | 58003 | 9.7 | 378.2 | N | 6095 | 46.0 | 0.58 | 199.2 | 61.6 | -0.30 | 17.3 | 75 | I | Am |
| Ecuador | 16.44 | 103.02 | Me | 255.01 | 80681 | 14.6 | 2233.1 | N | 5746 | 44.1 | 0.62 | 57.1 | 61.8 | -0.69 | 5.1 | 59 | C | Am |
| Egypt | 6.71 | 48.65 | La | 1001.08 | 248000 | 83.1 | 4768.0 | N | 3157 | 47.6 | 0.53 | 83.0 | NA | -0.73 | 7.2 | 35 | C | Af |
| El Salvador | 3.67 | 0.14 | Me | 20.54 | 23282 | 7.2 | 212.1 | N | 3896 | 40.3 | 0.54 | 349.9 | 64.5 | -0.13 | 10.5 | 24 | C | Am |
| Equatorial Guinea | 19.04 | 19.52 | Sm | 26.67 | 17483 | 0.7 | 223.6 | N | 17015 | 46.1 | 0.43 | 24.4 | 41.3 | -1.28 | NA | 28 | C | Af |
| Eritrea | 4.84 | 33.66 | Sm | 122.54 | 1856 | 5.6 | 445.8 | N | 532 | NA | 0.27 | 46.1 | 28.1 | -1.46 | NA | 59 | C | Af |
| Estonia | 7.97 | 1.52 | Sm | 45.82 | 22906 | 1.3 | 450.5 | N | 17632 | 33.1 | 0.83 | 28.4 | 80.0 | 1.11 | 16.1 | 81 | C | Eu |
| Eswatini | 4.08 | 2.18 | Sm | 17.11 | 4111 | 1.1 | 81.0 | N | 3468 | 60.5 | 0.50 | 65.7 | 46.1 | -0.60 | 6.1 | 55 | C | Af |
| Ethiopia | 4.07 | 73.54 | Me | 1127.37 | 42451 | 85.2 | 449.1 | N | 513 | 28.6 | 0.29 | 75.6 | 40.8 | -0.93 | 5.7 | 59 | C | Af |
| Federated States of Micronesia | 0.00 | NA | Mi | 0.63 | 301 | 0.1 | 0.0 | N | 3040 | 47.2 | 0.63 | 169.5 | 43.8 | 0.13 | NA | NA | I | Oc |
| Fiji | 2.23 | 2.05 | Sm | 18.93 | 3844 | 0.9 | 56.5 | N | 4645 | 43.0 | 0.72 | 49.9 | 55.7 | -0.40 | 40.6 | 104 | I | Oc |
| Finland | 9.16 | 29.31 | Me | 333.06 | 258000 | 5.3 | 3467.6 | Y | 47112 | 25.6 | 0.80 | 15.8 | 85.5 | 1.82 | 8.8 | 92 | C | Eu |
| France | 1.60 | 9.24 | La | 547.84 | 2700000 | 64.1 | 61724.0 | Y | 40606 | 29.5 | 0.79 | 116.9 | 86.6 | 1.19 | 8.9 | 71 | C | Eu |
| Gabon | 12.92 | 27.87 | Me | 259.97 | 15398 | 1.5 | 227.1 | N | 9026 | 36.7 | 0.58 | 5.8 | 51.7 | -0.58 | 3.1 | 56 | C | Af |
| Gambia | 0.21 | 0.10 | Sm | 10.50 | 317 | 1.8 | 4.0 | N | 490 | 40.5 | 0.30 | 169.8 | 52.4 | -0.59 | 21.9 | 50 | C | Af |
| Georgia | 8.11 | 7.59 | Me | 69.57 | 13668 | 4.6 | 592.7 | N | 3881 | 41.2 | 0.77 | 66.3 | 66.3 | 0.12 | 27.1 | 106 | C | As |
| Germany | 5.05 | 34.50 | La | 357.67 | 3580000 | 82.3 | 45222.2 | Y | 44375 | 28.9 | 0.86 | 230.2 | 86.7 | 1.49 | 10.8 | 97 | C | Eu |
| Ghana | 14.07 | 45.22 | Me | 238.67 | 35960 | 23.8 | 188.8 | N | 1520 | 38.3 | 0.49 | 99.9 | 57.6 | 0.06 | 7.1 | 110 | C | Af |
| Greece | 9.91 | 2.04 | Me | 131.35 | 270000 | 10.7 | 7104.0 | Y | 22102 | 33.1 | 0.76 | 81.7 | 79.4 | 0.38 | 18.6 | 80 | C | Eu |
| Grenada | 9.33 | 0.17 | Mi | 0.35 | 851 | 0.1 | 0.0 | N | 8294 | 42.9 | 0.72 | 261.3 | 53.6 | 0.38 | 20.2 | 108 | I | Am |
| Guatemala | 11.28 | 11.77 | Me | 108.81 | 49547 | 13.3 | 209.2 | N | 3503 | 43.4 | 0.42 | 122.0 | 63.2 | -0.59 | 8.2 | 63 | C | Am |
| Guinea | 0.43 | 2.80 | Me | 244.30 | 7513 | 10.1 | 170.0 | N | 680 | 33.3 | 0.26 | 41.2 | 45.1 | -1.19 | 5.4 | 82 | C | Af |
| Guinea Bissau | 19.81 | 13.40 | Sm | 32.83 | 963 | 1.5 | 18.6 | N | 614 | 40.2 | 0.34 | 46.7 | 38.5 | -1.12 | NA | 38 | C | Af |
| Guyana | 0.29 | 6.14 | Sm | 211.21 | 2620 | 0.8 | 35.7 | N | 3849 | 52.3 | 0.56 | 3.7 | NA | -0.36 | 7.3 | 89 | C | Am |
| Haiti | 0.00 | 1.15 | Sm | 26.89 | 7502 | 9.0 | 6.4 | N | 766 | 52.6 | 0.38 | 336.0 | 43.9 | -1.13 | 9.2 | 35 | I | Am |
| Honduras | 17.14 | 79.20 | Sm | 112.24 | 17335 | 7.8 | 215.4 | N | 2185 | 49.8 | 0.47 | 69.4 | 61.9 | -0.61 | 14.4 | 59 | C | Am |
| Hungary | 2.62 | 8.11 | Me | 93.20 | 135000 | 9.9 | 1409.0 | N | 13324 | 28.3 | 0.78 | 106.3 | 83.7 | 0.68 | 10.5 | 78 | C | Eu |
| Iceland | 15.29 | 138.92 | Me | 102.39 | 16426 | 0.3 | 23.7 | N | 49001 | 25.0 | 0.84 | 3.0 | 70.7 | 1.52 | 33.9 | 90 | I | Eu |
| India | 4.73 | 135.01 | VL | 3151.45 | 1730000 | 1166.1 | 44648.4 | Y | 1512 | 47.8 | 0.45 | 370.0 | 57.4 | -0.27 | 9.6 | 90 | C | As |
| Indonesia | 11.36 | 219.81 | VL | 1879.83 | 774000 | 240.3 | 5682.1 | N | 3493 | 43.8 | 0.55 | 127.8 | 61.8 | -0.38 | 6.2 | 99 | I | As |
| Iran | 3.38 | 154.25 | La | 1622.51 | 455000 | 66.4 | 11985.2 | N | 6250 | 39.4 | 0.61 | 40.9 | 49.5 | -1.07 | 7.7 | 48 | C | As |
| Iraq | 3.57 | 14.01 | Me | 437.37 | 169000 | 31.1 | 5027.9 | N | 5746 | 34.9 | 0.47 | 71.2 | 42.1 | -1.43 | 5.2 | 15 | C | As |
| Ireland | 0.87 | 0.34 | Me | 69.45 | 256000 | 4.2 | 1266.8 | N | 54862 | 30.1 | 0.84 | 60.5 | 82.0 | 1.48 | 5.8 | 33 | I | Eu |
| Israel | 20.40 | 10.27 | Me | 21.90 | 257000 | 7.2 | 15619.4 | Y | 34794 | 36.9 | 0.83 | 330.3 | 77.7 | 0.64 | 6.8 | 54 | C | As |
| Italy | 9.11 | 18.24 | La | 301.18 | 2120000 | 58.1 | 34069.0 | Y | 34373 | 32.8 | 0.75 | 193.0 | 82.1 | 0.53 | 11.1 | 96 | C | Eu |
| Jamaica | 4.92 | 0.07 | Sm | 11.03 | 13739 | 2.8 | 119.7 | N | 4954 | 55.9 | 0.64 | 256.1 | 65.9 | 0.05 | 30.3 | 39 | I | Am |
| Japan | 13.12 | 22.68 | La | 373.51 | 5220000 | 127.1 | 49562.4 | Y | 41886 | 30.3 | 0.79 | 340.2 | 74.4 | 1.27 | 7.4 | 98 | I | As |
| Jordan | 0.86 | 2.92 | Me | 88.86 | 29470 | 6.3 | 1547.0 | N | 3943 | 39.2 | 0.67 | 71.4 | 75.1 | -0.07 | 19.4 | 43 | C | As |
| Kazakhstan | 1.81 | 41.18 | La | 2714.27 | 168000 | 15.4 | 1780.4 | N | 11145 | 28.7 | 0.75 | 5.7 | 55.0 | -0.53 | 6.2 | 92 | C | As |
| Kenya | 8.84 | 135.12 | Me | 585.70 | 48808 | 39.0 | 733.5 | N | 1211 | 42.1 | 0.47 | 66.6 | 56.4 | -0.67 | 9.8 | 86 | C | Af |
| Kiribati | 22.35 | 0.39 | Mi | 1.03 | 164 | 0.1 | 0.0 | N | 1626 | 43.3 | 0.59 | 109.2 | 43.1 | 0.06 | 21.8 | 43 | I | Oc |
| Kuwait | 13.06 | 10.11 | Me | 17.47 | 60449 | 2.7 | 2346.7 | N | 40842 | 36.3 | 0.58 | 154.0 | 67.7 | 0.01 | 5.4 | 31 | C | As |
| Kyrgyzstan | 6.33 | 63.15 | Me | 198.99 | 5926 | 5.4 | 194.2 | N | 1135 | 32.4 | 0.68 | 27.3 | 58.9 | -0.84 | 3.9 | 70 | C | As |
| Laos | 0.00 | NA | Me | 228.11 | 9697 | 6.8 | 23.7 | N | 1772 | 40.2 | 0.41 | 30.0 | 39.7 | -0.87 | 14.2 | NA | C | As |
| Latvia | 9.36 | 9.16 | Me | 64.58 | 28890 | 2.2 | 347.6 | N | 13914 | 37.1 | 0.78 | 34.6 | 73.1 | 0.70 | 9.0 | 106 | C | Eu |
| Lebanon | 2.40 | 0.35 | Me | 10.00 | 40455 | 4.0 | 1725.2 | N | 8640 | 39.0 | 0.65 | 401.7 | 68.6 | -0.72 | 19.4 | 75 | C | As |
| Lesotho | 21.31 | 0.70 | Sm | 30.11 | 2335 | 2.1 | 47.8 | N | 1201 | 49.5 | 0.48 | 70.8 | 51.6 | -0.18 | 12.2 | 48 | C | Af |
| Liberia | 13.40 | 16.57 | Sm | 95.30 | 1541 | 3.4 | 10.3 | N | 420 | 32.4 | 0.41 | 36.1 | 52.5 | -0.81 | NA | 35 | C | Af |
| Libya | 0.35 | 40.00 | Me | 1623.76 | 65432 | 6.3 | 722.7 | N | 8862 | 29.1 | 0.61 | 3.9 | 53.8 | -1.33 | 7.1 | 40 | C | Af |
| Liechtenstein | 14.05 | 0.09 | Mi | 0.14 | 5534 | 0.0 | 0.0 | N | 161621 | NA | 0.80 | 253.3 | NA | 1.57 | 4.7 | 57 | C | Eu |
| Lithuania | 5.22 | 5.84 | Me | 64.94 | 42769 | 3.6 | 428.9 | N | 14583 | 33.4 | 0.82 | 54.7 | 75.6 | 0.81 | 5.3 | 58 | C | Eu |
| Luxembourg | 24.56 | 3.06 | Sm | 2.61 | 57249 | 0.5 | 263.4 | N | 108907 | 27.9 | 0.74 | 188.5 | 84.3 | 1.71 | 5.1 | 53 | C | Eu |
| Madagascar | 2.40 | 20.99 | Me | 592.98 | 9487 | 20.7 | 72.5 | N | 433 | 37.8 | 0.47 | 34.8 | 44.2 | -0.66 | 13.7 | 91 | I | Af |
| Malawi | 13.70 | 31.14 | Me | 119.40 | 6032 | 14.3 | 49.3 | N | 385 | 39.4 | 0.40 | 119.5 | 40.4 | -0.35 | 7.2 | 105 | C | Af |
| Malaysia | 9.51 | 45.42 | La | 327.88 | 275000 | 25.7 | 4494.6 | N | 10211 | 43.3 | 0.64 | 78.4 | 78.5 | 0.32 | 13.7 | 114 | I | As |
| Maldives | 3.38 | 0.01 | Mi | 0.11 | 2996 | 0.4 | 92.6 | N | 8489 | 45.4 | 0.49 | 3644.5 | 49.2 | -0.34 | 79.4 | 23 | I | As |
| Mali | 3.96 | 287.53 | Me | 1252.72 | 11895 | 12.7 | 186.9 | N | 786 | 33.8 | 0.24 | 10.1 | NA | -0.59 | 10.2 | 68 | C | Af |
| Malta | 20.92 | 0.23 | Mi | 0.33 | 9548 | 0.4 | 55.4 | N | 23581 | 28.0 | 0.71 | 1244.1 | 77.4 | 1.16 | 26.7 | 85 | I | Eu |
| Marshall Islands | 4.90 | 0.09 | Sm | 0.16 | 173 | 0.1 | 0.0 | N | 3442 | NA | NA | 391.5 | 44.5 | -0.09 | NA | 16 | I | Oc |
| Mauritania | 1.15 | 54.73 | Sm | 1036.39 | 4645 | 3.1 | 131.5 | N | 1285 | 32.8 | 0.32 | 3.0 | 49.7 | -0.83 | NA | 40 | C | Af |
| Mauritius | 3.69 | 0.56 | Me | 2.01 | 10925 | 1.3 | 17.7 | N | 9310 | 34.9 | 0.64 | 637.5 | 69.1 | 0.81 | 25.6 | 67 | I | Af |
| Mexico | 2.45 | 36.39 | VL | 1957.85 | 1120000 | 111.2 | 6373.1 | N | 9533 | 46.4 | 0.60 | 56.8 | 64.3 | -0.18 | 16.0 | 101 | C | Am |
| Moldova | 0.72 | 0.60 | Sm | 33.21 | 6524 | 4.3 | 26.3 | N | 1981 | 34.7 | 0.68 | 130.1 | 64.2 | -0.36 | 3.4 | 47 | C | Eu |
| Mongolia | 15.26 | 534.65 | Me | 1564.66 | 9208 | 3.0 | 85.9 | N | 3857 | 37.9 | 0.66 | 1.9 | 59.5 | -0.14 | 9.4 | 53 | C | As |
| Montenegro | 0.39 | 0.64 | Sm | 13.73 | 4262 | 0.7 | 70.4 | N | 6956 | 30.9 | 0.78 | 49.0 | 66.6 | 0.09 | 22.1 | 66 | C | Eu |
| Morocco | 1.30 | 15.55 | La | 591.74 | 97900 | 34.9 | 3289.0 | Y | 2961 | 36.3 | 0.42 | 58.9 | 65.6 | -0.31 | 18.5 | 76 | C | Af |
| Mozambique | 6.42 | 228.60 | Me | 788.45 | 12839 | 21.7 | 116.3 | N | 522 | 40.4 | 0.30 | 27.5 | 49.0 | -0.42 | 9.0 | 75 | C | Af |
| Myanmar | 5.53 | 118.36 | La | 663.16 | 50704 | 48.1 | 1924.2 | N | 1159 | 37.6 | 0.36 | 72.6 | 32.7 | -1.44 | 6.6 | 98 | C | As |
| Namibia | 16.11 | 509.85 | Me | 822.71 | 11083 | 2.1 | 393.2 | N | 5230 | 56.1 | 0.53 | 2.6 | 58.1 | 0.35 | 14.9 | 52 | C | Af |
| Nauru | 0.00 | NA | Mi | 0.02 | 76 | 0.0 | 0.0 | N | 8290 | 52.0 | NA | 658.2 | NA | 0.02 | NA | NA | I | Oc |
| Nepal | 8.41 | 36.11 | Me | 147.11 | 17131 | 28.6 | 266.6 | N | 691 | 41.1 | 0.41 | 194.2 | 41.1 | -0.82 | 7.5 | 45 | C | As |
| Netherlands | 12.33 | 11.31 | Me | 37.10 | 847000 | 16.7 | 10751.8 | Y | 49643 | 26.5 | 0.86 | 450.5 | 88.9 | 1.67 | 5.2 | 88 | C | Eu |
| New Zealand | 27.31 | 125.85 | Me | 268.49 | 163000 | 4.2 | 1972.7 | Y | 39586 | 32.2 | 0.89 | 15.7 | 78.2 | 1.80 | 17.5 | 178 | I | Oc |
| Nicaragua | 9.99 | 38.16 | Me | 128.69 | 10213 | 5.9 | 60.5 | N | 1867 | 42.8 | 0.49 | 45.8 | 59.6 | -0.58 | 10.7 | 39 | C | Am |
| Niger | 15.90 | 979.24 | Me | 1181.30 | 6481 | 15.3 | 94.3 | N | 384 | 32.5 | 0.16 | 13.0 | 44.1 | -0.65 | 4.9 | 64 | C | Af |
| Nigeria | 11.65 | 59.06 | La | 907.50 | 375000 | 149.2 | 2104.2 | N | 2666 | 39.0 | 0.46 | 164.4 | 55.4 | -1.12 | 4.7 | 106 | C | Af |
| Niue | 22.02 | 0.04 | Sm | 0.26 | 10 | 0.0 | 0.0 | N | NA | NA | NA | 5.3 | NA | NA | NA | 20 | I | Oc |
| North Korea | 2.58 | 13.20 | La | 122.38 | NA | 22.7 | 1000.0 | N | 696 | 31.0 | NA | 185.2 | NA | -1.58 | NA | 59 | C | As |
| North Macedonia | 8.06 | 7.36 | Sm | 25.39 | 10043 | 2.1 | 135.8 | N | 5008 | 36.2 | 0.66 | 81.4 | 67.0 | -0.07 | 6.7 | 70 | C | Eu |
| Norway | 12.54 | 34.21 | Me | 382.07 | 447000 | 4.7 | 6619.1 | Y | 90835 | 24.9 | 0.88 | 12.2 | 84.8 | 1.73 | 9.1 | 101 | C | Eu |
| Oman | 2.56 | 44.61 | Me | 311.21 | 65075 | 3.4 | 7667.1 | N | 19391 | NA | 0.61 | 11.0 | 62.0 | 0.21 | 7.3 | 24 | C | As |
| Pakistan | 10.33 | 106.29 | La | 872.94 | 213000 | 176.2 | 7363.4 | Y | 1284 | 36.0 | 0.33 | 201.9 | 53.3 | -1.09 | 6.9 | 63 | C | As |
| Palau | 0.00 | NA | Sm | 0.49 | 226 | 0.0 | 0.0 | N | 11432 | 46.4 | 0.79 | 42.3 | 51.1 | 1.15 | NA | NA | I | Oc |
| Panama | 10.61 | 57.01 | Me | 74.53 | 37666 | 3.4 | 389.7 | N | 11270 | 46.8 | 0.65 | 45.1 | 69.3 | 0.13 | 16.2 | 58 | C | Am |
| Papua New Guinea | 2.82 | 0.84 | Me | 465.15 | 17191 | 6.1 | 73.2 | N | 2610 | 52.8 | 0.36 | 13.0 | 51.4 | -0.62 | 1.9 | 50 | I | Oc |
| Paraguay | 2.43 | 72.54 | Me | 399.90 | 23251 | 7.0 | 270.3 | N | 4064 | 45.7 | 0.57 | 17.5 | 61.6 | -0.60 | 4.8 | 63 | C | Am |
| Peru | 6.73 | 252.35 | La | 1289.87 | 164000 | 29.5 | 2213.8 | N | 6048 | 47.3 | 0.64 | 22.9 | NA | -0.22 | 10.1 | 116 | C | Am |
| Philippines | 13.64 | 116.76 | La | 293.24 | 232000 | 98.0 | 2840.8 | N | 2642 | 46.3 | 0.60 | 334.1 | 63.2 | -0.40 | 19.7 | 95 | I | As |
| Poland | 1.50 | 3.84 | La | 313.43 | 493000 | 38.5 | 9245.9 | N | 13248 | 31.6 | 0.80 | 122.8 | 76.4 | 0.79 | 4.5 | 86 | C | Eu |
| Portugal | 1.92 | 6.96 | Me | 91.39 | 231000 | 10.7 | 4399.3 | Y | 21300 | 34.9 | 0.69 | 117.2 | 81.4 | 1.00 | 10.6 | 47 | C | Eu |
| Qatar | 11.31 | 0.01 | Sm | 11.15 | 149000 | 0.8 | 2788.3 | N | 77950 | 42.9 | 0.64 | 74.7 | 71.0 | 0.56 | 10.1 | 39 | C | As |
| Republic of Congo | 11.38 | 136.95 | Me | 344.89 | 11461 | 4.0 | 388.2 | N | 2572 | 43.6 | 0.49 | 11.6 | 49.5 | -1.06 | 4.1 | 108 | C | Af |
| Romania | 2.34 | 6.17 | La | 236.38 | 183000 | 22.2 | 2486.3 | N | 9166 | 32.0 | 0.73 | 94.0 | NA | NA | 5.2 | 92 | C | Eu |
| Russia | 6.63 | 369.60 | VL | 16953.09 | 1700000 | 140.0 | 68121.0 | Y | 12541 | 41.9 | 0.77 | 8.3 | 67.0 | -0.74 | 5.0 | 199 | C | Eu |
| Rwanda | 8.89 | 10.27 | Sm | 25.31 | 6592 | 10.5 | 82.2 | N | 667 | 45.8 | 0.36 | 413.9 | 45.1 | -0.25 | 11.2 | 89 | C | Af |
| Saint Kitts and Nevis | 3.31 | 0.26 | Mi | 0.27 | 775 | 0.0 | 0.0 | N | 15075 | 41.3 | 0.65 | 148.1 | NA | 0.73 | 25.1 | 33 | I | Am |
| Saint Lucia | 15.71 | 0.92 | Mi | 0.61 | 1444 | 0.2 | 0.0 | N | 8635 | 44.9 | 0.65 | 264.8 | 54.4 | 0.76 | 39.6 | 102 | I | Am |
| Saint Vincent and the Grenadines | 34.20 | 0.50 | Mi | 0.37 | 704 | 0.1 | 0.0 | N | 6558 | 47.7 | 0.65 | 284.4 | 48.6 | 0.76 | 22.3 | 70 | I | Am |
| Samoa | 1.72 | 0.50 | Mi | 2.78 | 714 | 0.2 | 0.0 | N | 4030 | 48.7 | 0.67 | 79.1 | 50.6 | 0.41 | NA | 40 | I | Oc |
| San Marino | 0.00 | NA | Mi | 0.06 | 2047 | 0.0 | 0.0 | N | 57165 | NA | NA | 502.7 | NA | 1.14 | NA | NA | C | Eu |
| São Tomé and Príncipe | 29.96 | 2.53 | Mi | 1.04 | 252 | 0.2 | 2.4 | N | 1513 | 27.6 | 0.44 | 205.1 | 42.7 | -0.32 | 31.0 | 12 | I | Af |
| Saudi Arabia | 2.21 | 138.54 | La | 1921.73 | 610000 | 28.7 | 56042.6 | N | 22646 | 32.0 | 0.66 | 14.9 | 63.2 | -0.34 | 10.2 | 31 | C | As |
| Senegal | 16.50 | 82.83 | Me | 196.22 | 13758 | 13.7 | 219.8 | N | 1013 | 35.5 | 0.29 | 69.9 | 59.3 | -0.24 | 11.0 | 64 | C | Af |
| Serbia | 3.57 | 6.38 | Me | 77.57 | 42403 | 7.4 | 902.1 | N | 5816 | 33.8 | 0.71 | 95.1 | NA | -0.11 | 6.7 | 70 | C | Eu |
| Seychelles | 32.33 | 1.64 | Mi | 0.49 | 1172 | 0.1 | 13.7 | N | 13938 | 39.5 | 0.66 | 179.6 | 60.7 | 0.20 | 58.1 | 45 | I | Af |
| Sierra Leone | 5.24 | 7.40 | Sm | 71.61 | 3444 | 6.4 | 32.0 | N | 561 | 34.9 | 0.32 | 89.9 | 46.3 | -0.70 | 3.6 | 102 | C | Af |
| Singapore | 6.58 | 0.31 | Me | 0.51 | 257000 | 4.7 | 8733.5 | N | 53304 | 40.0 | 0.72 | 9123.5 | 82.1 | 1.53 | 9.9 | 28 | I | As |
| Slovakia | 8.34 | 7.60 | Me | 48.46 | 93337 | 5.5 | 1135.0 | N | 17365 | 25.6 | 0.78 | 112.7 | 79.9 | 0.75 | 6.2 | 123 | C | Eu |
| Slovenia | 5.42 | 8.40 | Me | 20.33 | 48541 | 2.0 | 604.9 | N | 23006 | 25.1 | 0.83 | 98.7 | 77.9 | 0.94 | 12.6 | 94 | C | Eu |
| Solomon Islands | 0.04 | 0.11 | Sm | 27.20 | 885 | 0.6 | 44.8 | N | 1799 | 47.1 | 0.42 | 21.9 | 37.3 | -0.42 | 9.0 | 64 | I | Oc |
| Somalia | 0.00 | NA | Me | 471.82 | 5785 | 9.8 | 51.0 | N | 421 | 36.2 | NA | 20.8 | NA | -2.27 | NA | NA | C | Af |
| South Africa | 5.96 | 39.91 | La | 1219.82 | 340000 | 49.1 | 3839.4 | N | 6770 | 58.2 | 0.66 | 40.2 | 67.2 | 0.25 | 9.3 | 115 | C | Af |
| South Korea | 0.01 | 26.50 | Me | 98.54 | 1210000 | 48.5 | 31440.7 | N | 25553 | 30.8 | 0.82 | 492.3 | 75.0 | 0.77 | 5.1 | 56 | C | As |
| South Sudan | 10.28 | 198.87 | Me | 626.86 | 13407 | 10.6 | 1010.4 | N | 1221 | 42.6 | 0.30 | 16.9 | NA | NA | 5.2 | 79 | C | Af |
| Spain | 4.13 | 11.51 | La | 506.88 | 1400000 | 40.5 | 18450.8 | Y | 28914 | 33.6 | 0.76 | 79.9 | 82.6 | 0.86 | 14.2 | 64 | C | Eu |
| Sri Lanka | 31.41 | 13.14 | Me | 66.29 | 62119 | 21.3 | 1720.2 | N | 3500 | 52.0 | 0.71 | 321.7 | 57.7 | -0.33 | 11.4 | 143 | I | As |
| Sudan | 0.45 | 84.51 | Me | 1857.64 | 70162 | 25.9 | 3140.1 | N | 2014 | 33.2 | 0.27 | 14.0 | 42.0 | -1.59 | 5.2 | 83 | C | Af |
| Suriname | 11.28 | 116.69 | Sm | 145.12 | 4261 | 0.5 | 27.8 | N | 8506 | 54.6 | 0.61 | 3.3 | 45.6 | -0.09 | 2.7 | 57 | C | Am |
| Sweden | 10.67 | 55.27 | Me | 446.17 | 519000 | 9.1 | 5982.5 | N | 55861 | 25.3 | 0.83 | 20.3 | 87.3 | 1.77 | 9.6 | 109 | C | Eu |
| Switzerland | 7.31 | 1.70 | Me | 41.44 | 627000 | 7.6 | 4436.9 | N | 82883 | 29.4 | 0.84 | 183.5 | 88.2 | 1.74 | 9.6 | 104 | C | Eu |
| Syria | 0.00 | NA | La | 185.94 | 40405 | 20.2 | 1656.8 | N | 2058 | 40.1 | 0.48 | 108.5 | 47.7 | -1.39 | 13.0 | NA | C | As |
| Tajikistan | 19.58 | 233.58 | Me | 142.24 | 6621 | 7.3 | 69.7 | N | 912 | 43.2 | 0.65 | 51.7 | 45.2 | -1.13 | 8.2 | 80 | C | As |
| Tanzania | 27.86 | 415.74 | Me | 941.51 | 36732 | 41.0 | 342.7 | N | 837 | 43.6 | 0.38 | 43.6 | 48.6 | -0.40 | 13.3 | 113 | C | Af |
| Thailand | 18.82 | 37.16 | La | 514.45 | 358000 | 65.9 | 5222.4 | N | 5752 | 44.1 | 0.57 | 128.1 | 67.3 | -0.30 | 20.6 | 60 | C | As |
| Timor-Leste | 9.38 | 6.75 | Sm | 15.08 | 1136 | 1.1 | 32.3 | N | 1122 | 37.2 | 0.47 | 75.0 | NA | NA | NA | 18 | I | As |
| Togo | 10.30 | 21.67 | Sm | 56.86 | 3670 | 6.0 | 64.2 | N | 563 | 38.7 | 0.44 | 105.9 | 53.2 | -0.85 | 8.3 | 79 | C | Af |
| Tonga | 2.89 | 0.13 | Mi | 0.60 | 396 | 0.1 | 6.8 | N | 4059 | 42.0 | 0.73 | 200.5 | 44.7 | -0.06 | 18.3 | 39 | I | Oc |
| Trinidad and Tobago | 29.30 | 5.37 | Sm | 5.12 | 24004 | 1.3 | 171.7 | N | 18169 | 41.7 | 0.68 | 255.7 | 62.5 | 0.15 | 8.4 | 90 | I | Am |
| Tunisia | 0.25 | 1.66 | Me | 156.61 | 44118 | 10.5 | 721.8 | N | 4074 | 34.4 | 0.58 | 67.0 | 65.2 | -0.21 | 13.7 | 70 | C | Af |
| Turkey | 0.00 | NA | La | 780.08 | 817000 | 76.8 | 17101.8 | Y | 11464 | 40.9 | 0.58 | 98.5 | 67.5 | -0.12 | 12.5 | NA | C | As |
| Turkmenistan | 3.45 | 52.37 | Me | 470.85 | 29383 | 4.9 | 787.5 | N | 6408 | 35.4 | 0.63 | 10.4 | 38.6 | -1.37 | NA | 90 | C | As |
| Tuvalu | 5.18 | 0.00 | Mi | 0.02 | 34 | 0.0 | 0.0 | N | 3350 | 44.3 | NA | 537.5 | NA | 0.15 | NA | 15 | I | Oc |
| Uganda | 11.93 | 39.04 | Me | 241.85 | 21123 | 32.4 | 449.2 | N | 631 | 38.6 | 0.42 | 133.8 | 52.6 | -0.59 | 1.8 | 89 | C | Af |
| Ukraine | 2.50 | 25.90 | La | 599.09 | 142000 | 45.7 | 4092.4 | N | 3119 | 25.5 | 0.76 | 76.3 | 69.6 | -0.62 | 5.6 | 97 | C | Eu |
| United Arab Emirates | 0.04 | 0.27 | Me | 71.09 | 334000 | 4.8 | 16992.0 | Y | 40307 | 44.0 | 0.64 | 67.5 | 73.3 | 0.55 | 12.1 | 22 | C | As |
| United Kingdom | 10.60 | 6.19 | La | 243.78 | 2740000 | 62.3 | 58692.1 | Y | 42322 | 33.4 | 0.85 | 255.4 | 87.6 | 1.42 | 10.8 | 119 | I | Eu |
| United States of America | 9.40 | 947.29 | VL | 9833.52 | 16100000 | 314.0 | 646000.0 | Y | 53015 | 37.5 | 0.88 | 31.9 | 79.8 | 1.26 | 8.1 | 146 | C | Am |
| Uruguay | 0.18 | 1.69 | Me | 177.34 | 44541 | 3.5 | 809.2 | N | 15080 | 39.0 | 0.68 | 19.7 | NA | 0.82 | 9.6 | 41 | C | Am |
| Uzbekistan | 2.00 | 63.37 | Me | 447.83 | 47750 | 27.6 | 532.3 | N | 1841 | 35.2 | 0.70 | 61.6 | 42.4 | -1.25 | 3.1 | 92 | C | As |
| Vanuatu | 3.28 | 0.02 | Sm | 12.30 | 715 | 0.2 | 0.0 | N | 3055 | 44.1 | 0.52 | 17.8 | 48.2 | 0.18 | 44.5 | 35 | I | Oc |
| Venezuela | 18.90 | 513.52 | La | 912.68 | 353000 | 26.8 | 4132.3 | N | 12994 | 37.2 | 0.62 | 29.4 | 55.8 | -1.31 | 9.0 | 81 | C | Am |
| Vietnam | 7.51 | 12.23 | La | 328.89 | 145000 | 87.0 | 3243.9 | N | 1845 | 41.2 | 0.54 | 264.4 | 54.4 | -0.50 | 9.1 | 56 | C | As |
| Yemen | 0.00 | NA | Me | 453.07 | 32142 | 23.8 | 1493.9 | N | 1386 | 41.1 | 0.31 | 52.6 | 53.6 | -1.37 | 5.7 | NA | C | As |
| Zambia | 17.65 | 223.14 | Me | 751.91 | 21394 | 11.9 | 318.2 | N | 1574 | 50.0 | 0.52 | 15.8 | 53.5 | -0.28 | 7.0 | 84 | C | Af |
| Zimbabwe | 17.12 | 147.76 | Me | 389.34 | 11908 | 12.6 | 237.0 | N | 950 | 41.0 | 0.51 | 32.4 | 50.5 | -1.43 | 8.1 | 92 | C | Af |
